# Acquired *RAD51C* promoter methylation loss causes PARP inhibitor resistance in high grade serous ovarian carcinoma

**DOI:** 10.1101/2020.12.10.419176

**Authors:** Ksenija Nesic, Olga Kondrashova, Rachel M. Hurley, Cordelia McGehee, Cassandra J Vandenberg, Gwo-Yaw Ho, Elizabeth Lieschke, Genevieve Dall, Nirashaa Bound, Kristy Shield-Artin, Marc Radke, Ashan Musafer, Zi Qing Chai, Mohammad Reza Eftekhariyan Ghamsari, Maria I. Harrell, Damien Kee, Inger Olesen, Orla McNally, Nadia Traficante, Australian Ovarian Cancer Study, Anna DeFazio, David D. Bowtell, Elizabeth M. Swisher, S. John Weroha, Katia Nones, Nicola Waddell, Scott H. Kaufmann, Alexander Dobrovic, Matthew J. Wakefield, Clare L. Scott, Australian Ovarian Cancer Study (AOCS)

## Abstract

While loss of *BRCA1* promoter methylation has been shown to cause PARP inhibitor (PARPi) resistance in high-grade serous ovarian carcinoma (HGSC), the impacts of *RAD51C* methylation (me*RAD51C*) remain unresolved. In this study, three PARPi-responsive HGSC patient-derived xenografts (PDX) with *RAD51C* gene silencing and homologous recombination deficiency were found to have either homogeneous or heterogeneous patterns of me*RAD51C*. PDX could lose me*RAD51C* following PARPi treatment (rucaparib/niraparib), where a single unmethylated *RAD51C* copy was sufficient to drive PARPi-resistance. Genomic profiling revealed this resistance was acquired independently in two distinct PDX lineages. Furthermore, we describe a patient sample where 1/3 *RAD51C* gene copies lost methylation following neoadjuvant chemotherapy. We show me*RAD51C* is a positive predictive biomarker for PARPi response and should be screened for routinely in patients. However, methylation loss in a single gene copy is sufficient to cause PARPi resistance and should be carefully assessed in previously treated patients considering PARPi therapy.

## INTRODUCTION

Women diagnosed with high-grade serous ovarian carcinoma (HGSC) are generally responsive to standard-of-care platinum-based chemotherapy initially, and in those with inherited mutations in *BRCA1/2*, the addition of poly ADP ribose polymerase inhibitor inhibitors (PARPi) can significantly improve clinical outcomes ^1, 2^. However, the development of resistance is common and limits long-term effectiveness^3, 4^. While the impacts of deleterious homologous recombination repair (HRR) gene mutations (e.g. in *BRCA1*, *BRCA2* and *RAD51C*) on platinum and PARPi responses in HGSC are well recognized, consequences of HRR gene promoter methylation in these contexts have remained controversial^5, 6, 7, 8, 9^.

HRR gene methylation has been shown to correlate with reduced gene expression for the *BRCA1* and *RAD51C* genes, and is usually found to be mutually exclusive of deleterious *BRCA1/2* mutations^8, 9, 10, 11, 12, 13^. Pre-clinical studies have provided some evidence that silencing of the *BRCA1* and *RAD51C* genes by methylation results in HRR deficiency (HRD)^14, 15, 16^. Importantly, genomic scarring related to defective HRR and *BRCA1/2* mutations has also been detected in breast and ovarian cancers with *BRCA1* methylation (me*BRCA1*) and *RAD51C* methylation (me*RAD51C*)^12, 17, 18^. However, based on mixed outcomes in several PARPi clinical studies that included screening for HRR gene methylation, it has remained unclear whether me*BRCA1* and me*RAD51C* predicts PARPi response^8, 9, 10, 12, 13, 19^.

Our group recently determined, using a cohort of HGSC Patient Derived Xenograft (PDX) models, that the zygosity of me*BRCA1* is a key factor for determining PARPi and platinum response in HGSC^20^. This discovery was made possible by the fact that only human tumor cells are propagated as PDX, excluding normal stroma and other non-tumor human cells^21^, thus enabling accurate determination of zygosity in the tumors. Limitations of previous studies had included the use of non-quantitative methods, such as methylation-specific PCR (MSP)^8, 9^, as well as the presence of contaminating non-tumor tissue, which made it challenging to interpret the zygosity of me*BRCA1*^7^. Our study demonstrated that in a tumor, where multiple copies of *BRCA1* may be present, all copies must be methylated for PARPi response, and that losing methylation of a single *BRCA1* copy was sufficient to restore HRR DNA repair and cause PARPi resistance ^20^.

Although we are now beginning to better understand me*BRCA1* stability under platinum or PARPi treatment pressure, little is known about me*RAD51C* in HGSC. The RAD51C protein is critical for HRR DNA repair, playing a pivotal role in the resolution of Holliday junctions during recombination. *RAD51C* is a known ovarian cancer susceptibility gene^22, 23, 24^, and recently it was shown that deleterious mutations in this gene cause PARPi sensitivity, while secondary reversion mutations that restore function of at least a single *RAD51C* copy cause PARPi resistance^25^.

Me*RAD51C* is found in approximately 2% of HGSC^7, 19^, and its effects on PARPi response remain unclear^9^. This uncertainty is likely due to the same limitations encountered when studying me*BRCA1*, such as the ability of a single demethylated copy to cause therapy resistance^20^ and the difficulties in accurately assessing methylation zygosity (i.e. whether all copies of the gene are methylated) in the presence of contaminating stroma using qualitative or semi-quantitative assays ^20^.

We hypothesized that, as for me*BRCA1* HGSC^20^, me*RAD51C* zygosity could influence PARPi response in HGSC and that homozygous me*RAD51C* could become heterozygous under PARPi or platinum treatment pressure, thus leading to therapeutic resistance. Here we use several HGSC PDX models to characterize the impacts of me*RAD51C* on PARPi treatment and then to align these outcomes with those of a cohort of me*RAD51C* HGSC patients.

## RESULTS

### RAD51C methylation leads to gene silencing and PARPi response in HGSC PDX

PDX #183 was derived from chemo-naïve neoplastic cells obtained from a woman diagnosed with Stage III, HGSC. The PDX was confirmed to be HGSC at transplant 3 (T3) by histopathological and immunohistochemical (IHC) review (Supplementary Fig. 1). This model was found to be cisplatin sensitive (p < 0.001; Figure 1A; Supplementary Table 1), reflecting the patient’s response to first-line platinum chemotherapy (> 11 months from cessation of platinum chemotherapy until time of first progression). This PDX also demonstrated response to the PARP inhibitors, rucaparib and niraparib (Figure 1B-C), with a delay in median time to progression (TTP) of 8 vs 47 days for rucaparib 450 mg/kg (p<0.0001) and 8 vs 50 days for niraparib (p<0.0001) (Supplementary Table 1). A nonsense *TP53* mutation c.661G>T (p.E221*) was found by targeted sequencing using BROCA^26^, but no pathogenic HRR gene mutations were detected (Supplementary Table 2). A heterozygous missense *PARP1* mutation c.2008C>A (p.Q670K) in the regulatory domain was also detected in this PDX, but the effects of this mutation on PARPi activity are unclear ^27^. Given the observed *in vivo* PARPi response, we tested this model for both me*BRCA1* and me*RAD51C*. Me*BRCA1* was not detected in this model (data not shown). Me*RAD51C* was detected initially by a non-quantitative method, MSP (Figure 1D), and was confirmed to be homozygous using MS-HRM (Figure 1E). PDX #183 had no detectable *RAD51C* gene expression by RNA sequencing (Figure 1F) or protein expression by immunoblotting (Figure 1G) when compared with *RAD51C* wild-type controls PDX 13 and cell line PEO4, confirming that the homozygous me*RAD51C* was, indeed, causing gene silencing and thus likely causing the susceptibility to PARPi in this PDX model. Thus, PDX #183 was a cisplatin-sensitive and PARPi-responsive HGSC PDX model with me*RAD51C* and *RAD51C* gene silencing and consequent protein loss.

**Table 1.** Summary of clinical information and *meRAD51C* status of the patient cohort. AOCS-143 cellularity was based on a second section taken from the same tumor. Extended patient clinical details are reported in Supplementary Table 7. N/A –– data not available; FU – Follow-up; DOD –; DOD – Died of disease; TFI – Treatment-Free Interval. *pattern based only on MS-HRM.

| Patient ID | Surgical outcome | Lines of therapy received |  |  | First platinum TFI | Primary platinum status | Survival | Neoplastic cellularity (%) | *RAD51C* copy number | Methylated alleles % | Adjusted *meRAD51C* | *meRAD51C* profile |
| --- | --- | --- | --- | --- | --- | --- | --- | --- | --- | --- | --- | --- |
|  |  | Total | Platinum | Prior to sample collection |  |  |  |  |  |  |  |  |
| LS-158 | <5mm | 1 | 1 | 0 | 87 months | Sensitive | 87 months at last FU, alive | 90 | 2 | 93 | 100% | Homogeneous |
| LS-215 | No residual disease | 3 | 3 | 0 | 22 months | Sensitive | 41 months, DOD | 56 | 2 | 72 | 100% | Homogeneous |
| LS-267 | Suboptimal debulk | 3 | 1 | 0 | 18 months | Sensitive | 24 months, DOD | 59 | 2 | 75 | 100% | Homogeneous |
| AOCS-106 | Macroscopic disease ≤1 cm | 2 | 1 | 1 | 4 months | Resistant (sample post-neoadjuvant chemotherapy) | 12 months, DOD | 92 | 3 | 55 | 58% | Homogeneous and heterozygous |
| LS-376 | No residual disease | 5 | 3 | 0 | 2 months | Resistant | 25 months, DOD | 96 | 3 | 89 | 92% | Homogeneous |
| LS-28 | Suboptimal debulk, >2cm | ≥1 | ≥1 | 0 | N/A | Refractory | 25 months, DOD | N/A | N/A | 72 | N/A | Homogeneous |
| AOCS-120 | Macroscopic disease ≤1 cm | 6 | 4 | 6 | 17 months | Sensitive (sample from resistant recurrence) | 76 months, DOD | 93 | 3 | 96 | 100% | Heterogeneous |
| AOCS-143 | Macroscopic disease ≤1 cm | 9 | 6 | 0 | 7 months | Sensitive | 69 months, DOD | 78 | 2 | 81 | 100% | Heterogeneous |
| LS-473 | Suboptimal debulk, 3-4cm | 6 | 3 | 1 | 6 months | Resistant (sample post-neoadjuvant chemotherapy) | 45 months, DOD | 79 | 4 | 86 | 97% | Heterogeneous |
| LS-472 | Suboptimal debulk, 1.5cm | 4 | 1 | 0 | N/A | Resistant | 26 months, DOD | 97 | 1 | 94 | 99% | Heterogeneous |
| LS-467 | No residual disease | N/A | N/A | 0 | N/A | N/A | N/A | 76 | 3 | 88 | 100% | Heterogeneous |
| SFRC-1307 | Suboptimal debulk | 0 | 0 | 0 | N/A | N/A | 3 months at last FU, alive | N/A | N/A | N/A | N/A | Heterogeneous* |

**Figure 1.**
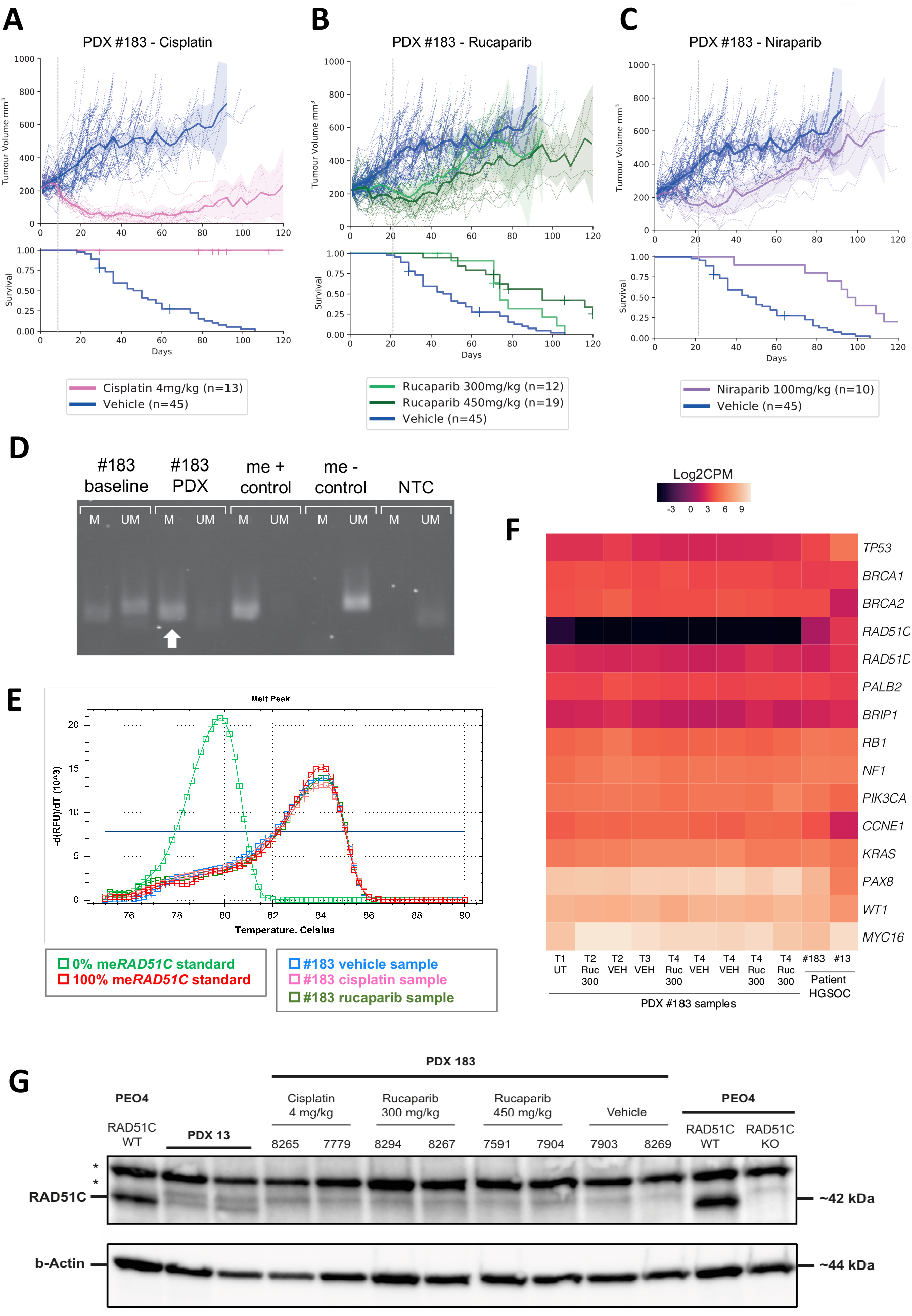
Characterization of a novel PARPi-responsive HGSC PDX with *RAD51C* promoter methylation. **(A)** PDX #183 response to cisplatin (p<0.0001, compared to DPBS), and **(B)** PARP inhibitors rucaparib (at 300 mg/kg and 450 mg/kg doses, p=0.01 and p<0.0001 respectively compared to DPBS) and **(C)** niraparib (p<0.0001, compared to DPBS) *in vivo*. See Supplementary Table 1 for all median TTH, TTP and p-values for survival comparison. Mean tumor volume (mm3) ± 95% CI (hashed lines are representing individual mice) and corresponding Kaplan–Meier survival analysis. Censored events are represented by crosses on Kaplan–Meier plot; n = individual mice. **(D)** MS-PCR of the *RAD51C* promoter in patient #183 (baseline) tumor and PDX #183 sample (arrow). “me +” – Positive me*RAD51C* control; “me –” – negative me*RAD51C* control; NTC – no n template control; M – methylated *RAD51C* reaction; UM – Unmethylated *RAD51C* reaction. **(E)** Homozygous me*RAD51C* detection in standard single-dose vehicle, cisplatin and rucaparib treated PDX samples by MS-HRM. Each line represents a measurements per tumor/sample. RFU – Relative fluorescence units, Y axis is the derivative of fluorescence over temperature (-d(RFU)/dT) versus temperature (T). **(F)** *RAD51C* mRNA expression by RNAseq (VEH: vehicle, RUC 300: rucaparib 300 mg/kg dose) compared to other genes frequently defective in HGSC. **(G)** RAD51C protein measured by immunoblotting in PDX #183 tumor samples following the indicated *in vivo* treatments. PEO4 cell line +/− RAD51C and PDX #13 (expressing RAD51C) were used as controls. *denotes non-specific bands.

PDX #1240 was derived from chemo-naïve neoplastic cells obtained from the ovary of a woman diagnosed with stage IIIC HGSC. The PDX was confirmed to be HGSC at T1 by histopathological and immunohistochemical (IHC) review (Supplementary Fig. 1). Like #183, this model was found to be highly cisplatin sensitive (p <0.0001) and responsive to PARPi rucaparib (p= 0.013; Supplementary Table 1; Supplementary Figure 2A). As the patient tumor sample was found to lack mutations in any core HRR genes (Supplementary Table 2), the PDX model was tested for HRR gene methylation, where homogeneous me*RAD51C* was detected on MS-HRM, while *BRCA1* methylation was absent (Supplementary Figure 2B-C).

### Heterogeneous RAD51C methylation patterns also cause gene silencing and PARPi response in HGSC PDX

The initial characterization of the PDX model PH039 has been published^28^. PH039 was re-established within our laboratory for further me*RAD51C* characterization and comparison with me*RAD51C* PDX #183.

Cryopreserved minced neoplastic material from the original PH039 PDX was transplanted into several NSG mice, and two PDX lineages derived from the initial transplant (T1) were used in this study – Lineage A (PH039-A) and Lineage B (PH039-B; Figure 2A; Supplementary Fig. 1). Although both of these lineages were responsive to PARPi, PH039-A lineage was consistently less responsive to PARPi compared with PH039-B. This was observed for treatments with rucaparib 450 mg/kg (TTP: PH039-A= 50 vs PH039-B= 71 days, *p*= 0.0004), and niraparib 100 mg/kg (TTP: PH039-A= 43 vs PH039-B= 64 days, *p*= 0.0766) (Figure 2B; Supplementary Fig. 3; Supplementary Table 3). Both lineages were maintained in order to study the basis of these differences. PH039-A and PH039-B were both cisplatin sensitive, though PH039-A had a more durable response (TTP = 134 days vs 106 days for PH039-B, *p*= 0.1269); Figure 2C; Supplementary Table 3).

**Figure 2.**
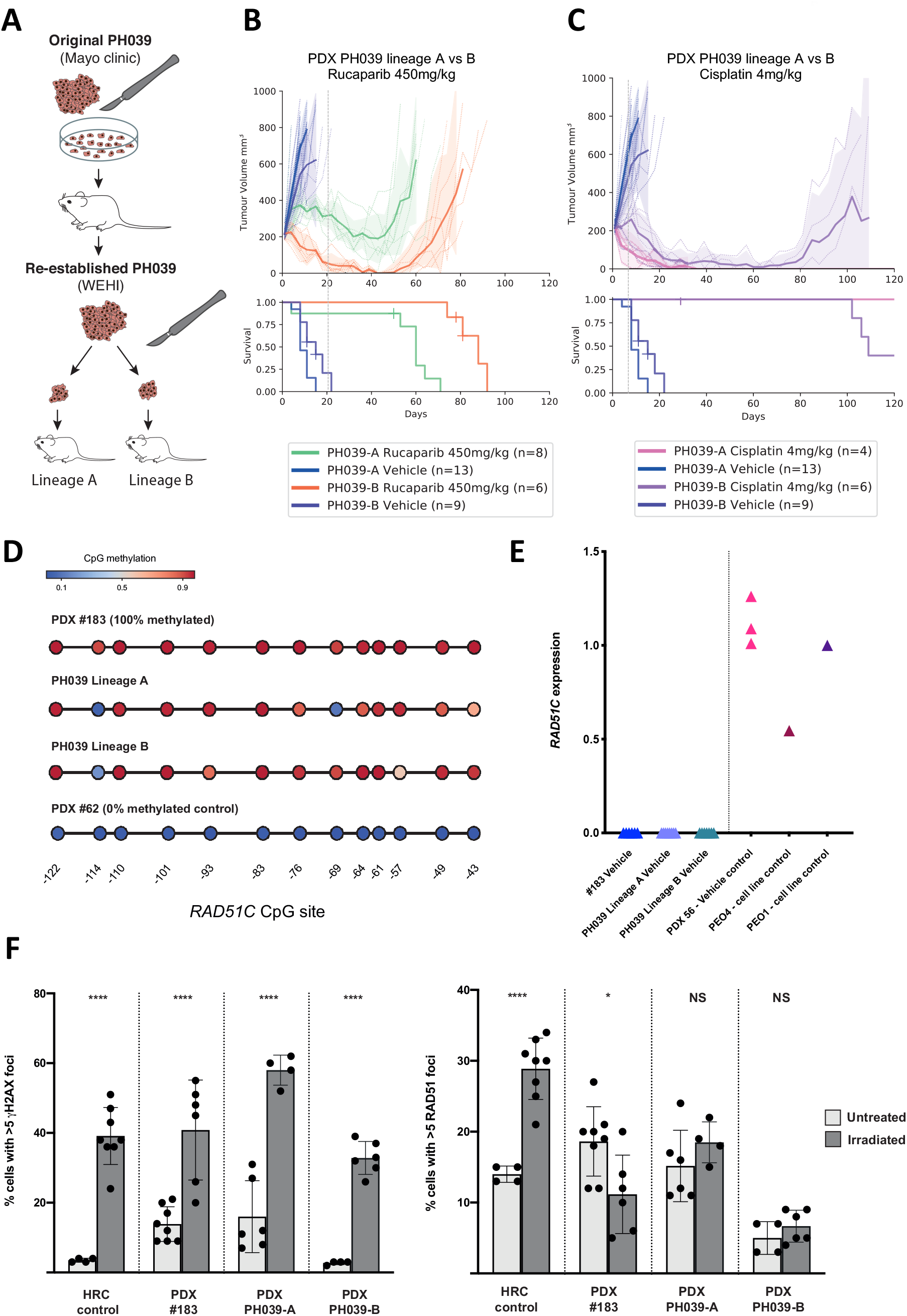
Heterogeneous *RAD51C* methylation patterns observed in PDX PH039 lineages still lead to *RAD51C* silencing, HRD and PARPi response *in vivo*. **(A)** PDX PH039 was re-established in our laboratory using cryopreserved tumor material sent to us from the Kaufmann group (Mayo Clinic). Two lineages of this PDX were formed following initial (T1)transplantation – Lineage A (PH039-A) and Lineage B (PH039-B). **(B)** While PH039 Lineage A was found to have some response to PARPi (SD; p<0.001; rucaparib shown here, niraparib shown in supplementary data), Lineage B was found to be more rucaparib-responsive (PR; p=0.067). **(C)** Both lineages were sensitive to cisplatin (Lineage A: CR; p>0.001 and lineage B: CR; p=0.01), although Lineage B tumors had an earlier progression timepoint compared to Lineage A. Mean tumor volume (mm3) ± 95% CI (dashed lines are individual mice) and corresponding Kaplan–Meier survival analysis. Censored events are represented by crosses on Kaplan–Meier plot; n = individual mice. **(D)** Targeted me*RAD51C* bisulfite sequencing revealed that vehicle-treated PH039 Lineage A tumors have reduced methylation of CpG sites −114 and −69 within the *RAD51C* promoter. While 50% of the Lineage B tumors assessed had a me*RAD51C* profile like that of Lineage A, the other 50% had a more methylated heterogeneous profile presented here (reduced methylation at CpG site −114). PDX #183, in contrast, is homogeneously methylated across the promoter. PDX #62 presented as me*RAD51C* negative control. **(E)** No vehicle-treated tumors from either of the PDX PH039 lineages had detectable *RAD51C* gene expression by qRT-PCR (fold change from *BRCA2*-mutant cell line PEO1).). HRR competent (HRC) PEO4 cell line and *BRCA1*-mutant PDX #56 vehicle samples included for additional controls. **(F)** PDX #183 and PH039-A and PH039-B tumors have limited RAD51 DNA repair foci-forming capacity relative to control HRC PDX control samples (PH038 and #931). Each data point represents a % cell count (geminin positive cells with ≥5 foci) from 1 of 2 individual tumor replicates. HRC control group consists of PDX models #931 and PH038 (50:50 # of tumors in irradiated and untreated groups). P-values calculated using Welch’s two-tailed t test in PRISM.. **** – p<0.0001; * – p<0.01; NS – not significant.

When assessing the me*RAD51C* patterns observed for each lineage, MS-HRM results for untreated PH039-A aliquots revealed a “heterogeneous” pattern of me*RAD51C* (causing a broad MS-HRM peak), consistent with independent findings for this model ^29^ (Supplementary Fig. 4A). Such heterogeneous methylation has been previously reported for other genes, including the *CDKN2B* gene^30^. It should be noted that this heterogeneous methylation is different to a heterozygous me*RAD51C* pattern, which would consist of two melt peaks – one methylated and one unmethylated (example in Supplementary Fig. 4B). Instead, a broad MS-HRM me*RAD51C* peak is characteristic of a highly variable mixture of epialleles (i.e. me*RAD51C* copies) containing different patterns of CpG methylation which then can form multiple heteroduplexes resulting in the broadened peak. Half of the tested vehicle-treated PH039-B aliquots matched this PH039-A profile; the other half, however, displayed a more-methylated heterogeneous profile – designated “right-shifted heterogeneous” (Supplementary Table 4).

To elucidate the underlying assortment of cancer epialleles corresponding to these me*RAD51C* MS-HRM profiles in PH039 (“heterogeneous” versus “right-shifted heterogeneous”), targeted me*RAD51C* bisulfite sequencing was performed on PDX aliquots with each profile. This analysis enables characterization of CpG methylation patterns for individual epialleles and also shows the frequency of each unique epiallele within the neoplastic sample. Sequencing revealed that the heterogeneous me*RAD51C* patterns in PH039-A represented a mixture of multiple epialleles, with a majority of these having reduced methylation of CpG sites at positions −114 and −69 (condensed amplicon summary in Figure 2D, full profiles in Supplementary Fig. 4C-G). However, it was found that the CpG site at position −69 was, in contrast, highly methylated in the vehicle-treated PH039-B samples with “right-shifted heterogeneous” MS-HRM profiles, thereby explaining the shifts in MS-HRM curves (Figure 2D; Supplementary Fig. 4D-G). Consistent with the MS-HRM data, PDX #183 tumor aliquots had highly methylated and homogeneous me*RAD51C* sequencing profiles relative to me*RAD51C* PH039 tumor aliquots (Figure 2D; Supplementary Fig. 5).

Interestingly, both the “heterogeneous” and “right-shifted heterogeneous” me*RAD51C* profiles of PH039-A and PH039-B were found to be associated with gene silencing in vehicle-treated samples, like the homogeneously methylated PDX #183 (Figure 2E). As a previous study of heterogeneous promoter methylation at the *CDKN2B* locus in acute leukemia found that epialleles with <40% CpG methylation across the promoter could re-express the gene (regardless of CpG position), we were interested in the minimum degree of methylation required in this genomic region for silencing to occur^31^. In our PDX samples, this was found to be 7 methylated CpG sites out of 13 sites in total (54% of region CpG sites methylated; Supplementary Fig. 5C).

Both PDX PH039 lineages and PDX #183 were also found to have impaired RAD51 foci forming capacity following DNA damage, confirming that homogeneous and heterogeneous me*RAD51C* patterns both cause *RAD51C* gene silencing and HRD (Figure 2F; Supplementary Fig. 6).

Given the complexity of the me*RAD51C* patterns uncovered by the targeted bisulfite amplicon assay in this study, we have summarized the definitions used to describe me*RAD51C* in Fig. 3. “Methylation zygosity” refers to the combinations of epialleles present within individual cells. These can be “homozygous methylated” (only highly/completely methylated epialleles present), “heterozygous methylated” (mixture of highly/completely methylated and unmethylated epialleles) and “unmethylated” (no methylated epialleles present). “Methylation pattern”, in contrast, describes epiallele diversity at the tumor/tissue level, with either “homogeneous” or “heterogeneous” mixtures of methylated epialleles detected.

**Figure 3.**
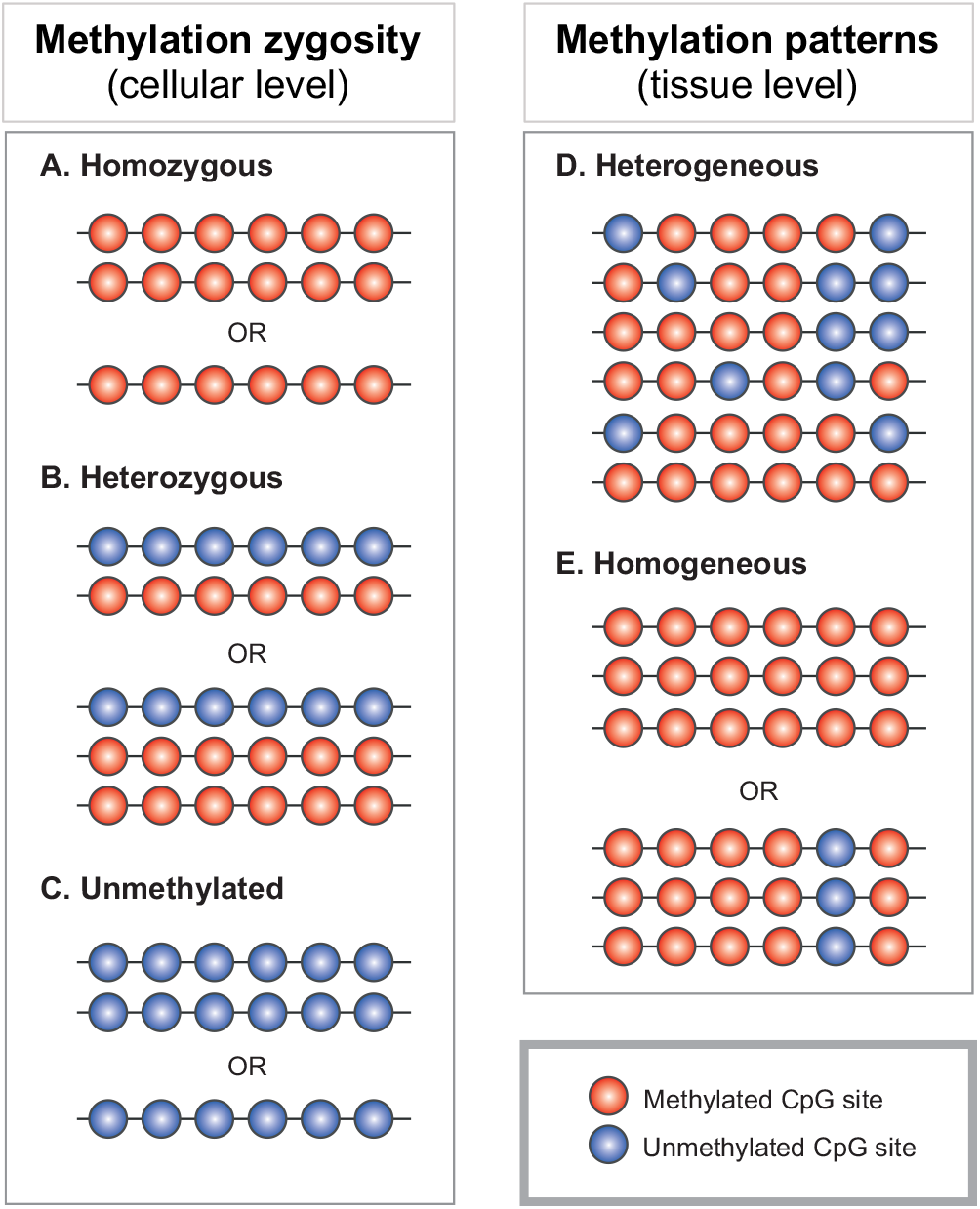
Definitions of promoter methylation states in tumors. The following definitions pertain to either methylation zygosity, which is describes combinations of epialleles present at the cellular level, and methylation patterns, which describe epiallele diversity at the tumor/tissue level. (A) “Homozygous methylation” describes when all epialleles in a cell are fully/highly methylated (all CpG sites in promoter are methylated) and the given gene is completely silenced. This includes cases where an allele has been lost due to aneuploidy and the remaining hemizygous allele is methylated. (B) “Heterozygous methylation” describes mixture of fully/highly methylated and unmethylated epialleles in each cell, where gene expression is permitted due to the presence of fully unmethylated epialleles. (C) Unmethylated cells have no methylated epialleles detected, and the gene is not silenced. (D) “Heterogeneous methylation” describes a highly heterogeneous mixture of epialleles with various CpG methylation patterns present in a tumor sample. This is represented as a broad peak on MS-HRM. (E) “Homogeneous methylation” describes a homogeneous mixture of highly methylated epialleles predominated by one particular CpG methylation pattern in a tumor sample. This can include samples with homozygous methylation, and is represented by a narrow peak on MS-HRM.

### Loss of RAD51C methylation results in RAD51C expression and PARPi resistance in HGSC PDX

In an effort to drive me*RAD51C* loss and PARPi resistance in PDX models, *in vivo* cyclical rucaparib re-treatment experiments were carried out. PDX #183 tumor aliquots were re-treated at a rucaparib dose of 300 mg/kg (Supplementary Fig. 7), while PH039-A and -B xenografts required a higher dose of 450 mg/kg due to their rapid growth rate (Figure 4A).

**Figure 4.**
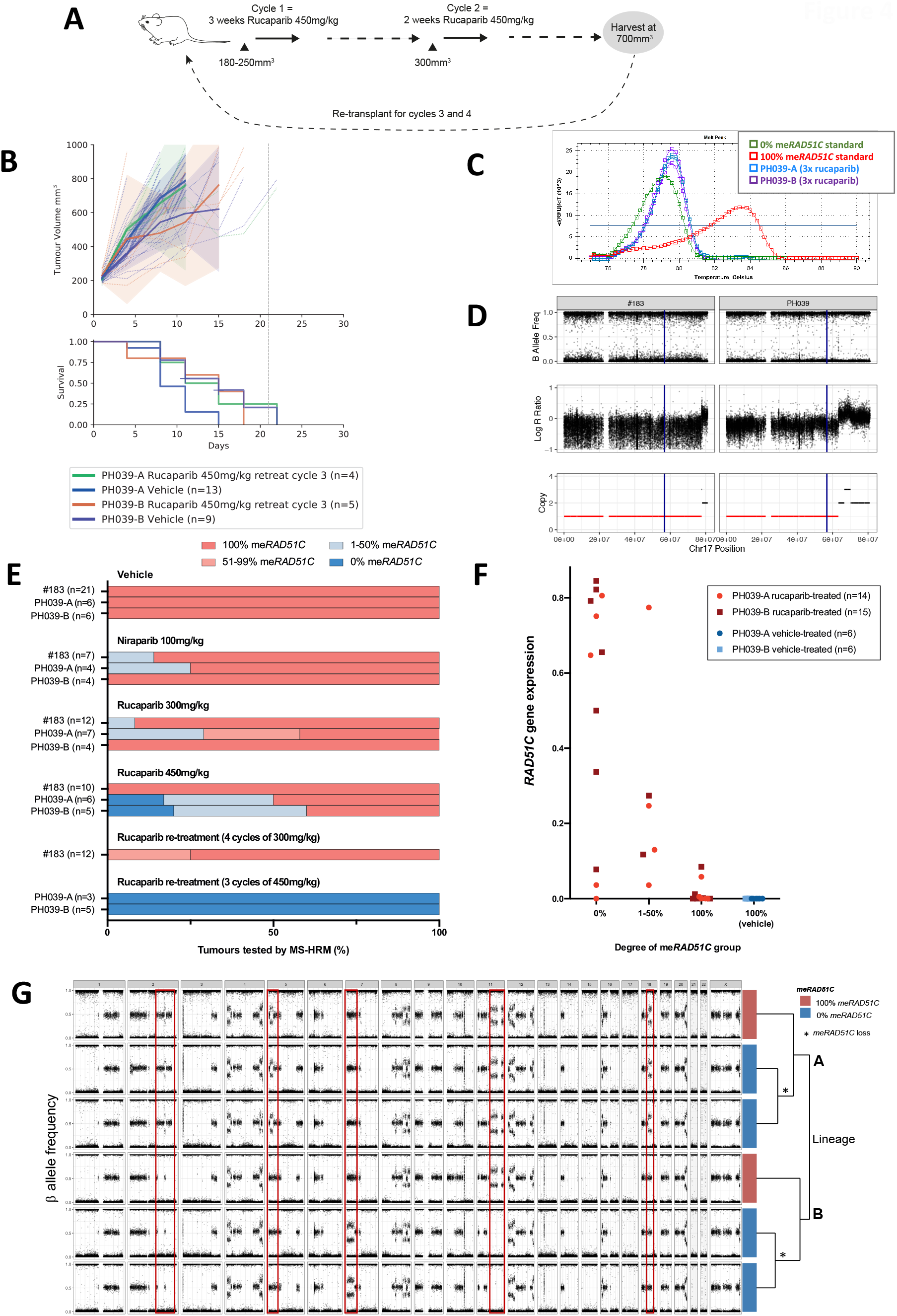
Rucaparib re-treatment of PDX PH039 lineages results in rucaparib-refractory tumors that have complete methylation loss at the *RAD51C* promoter. **(A)** PH039-A and PH039-B tumors were cyclically re-treated with rucaparib (two cycles per transplant) at a 450 mg/kg dose. **(B)** However, tumors of both PH039 lineages became so refractory to rucaparib after three cycles that none could receive a fourth treatment cycle (TTH, TTP and p values in Supplementary Table 3). Mean tumor volume (mm3) ± 95% CI (hashed lines are representing individual mice) and corresponding Kaplan– Meier survival analysis. Censored events are represented by crosses on Kaplan–Meier plot; n = individual mice. **(C)** All rucaparib-refractory cycle-3 PH039-A and PH039-B tumors tested had complete loss of me*RAD51C* by MS-HRM – two sample MS-HRM curves presented here. **(D)** Chr17 copy number profiles from SNP array data of PDX models revealed only one *RAD51C* gene copy (location indicated by blue vertical line) per cell in both #183 and PH039 Lineages A and B. **(E)** MS-HRM screening results for all PDX models revealed PH039-A to be most prone to me*RAD51C* loss following PARPi treatment. The most consistent complete loss of me*RAD51C* was observed in PH039-A and PH039-B tumors following three cycles of rucaparib. Presented is the % of tumors from each model with the degree of me*RAD51C* observed by MS-HRM screening following different PARPi or cisplatin treatments. Minimum of n=4 samples tested per treatment group. **(F)** Loss of me*RAD51C* measured by MS-HRM in rucaparib-treated (300 mg/kg,450 mg/kg and 450 mg/kg re-treatment) PH039 Lineage A and B tumor samples correlated with restored *RAD51C* gene expression by qRT-PCR (PH039-A R^2^ = 0.3279, p= 0.0323; PH039-B R^2^ = 0.6288, p= 0.0021). Weaker correlation between me*RAD51C* loss and *RAD51C* gene expression is possibly due to tumor heterogeneity, as different pieces of same PDX aliquot were assessed for me*RAD51C* and *RAD51C* expression. All vehicle-treated samples had 100% me*RAD51C* and undetectable levels of *RAD51C* gene expression, and are included here for reference. *RAD51C* expression normalized to that of PEO1 cell line. **(G)** SNP array analysis of post-rucaparib re-treatment PH039 lineage A and B tumor aliquots reveals copy number profile differences between lineages (large changes in red boxes). The dendrogram on the right of the figure was generated using the complete linkage method for hierarchical clustering, and presents the predicted pathway of tumor evolution.

Rucaparib re-treatment of PH039-A and PH039-B tumor aliquots resulted in recurrent cancers which were treatment-refractory by the third cycle of treatment and thus did not reach the fourth cycle (Figure 4B). All recurrent PH039-A and -B cancers were found to have complete loss of me*RAD51C* by the third cycle (Figure 4C). Interestingly, no heterozygous me*RAD51C* patterns were detected in these refractory cancers. This was explained by a somatic loss of heterozygosity of the chromosome 17, including the *RAD51C* locus, resulting in a single gene copy in both PDX PH039 and PDX #183 (before treatment; Figure 4D). Thus, heterozygous me*RAD51C*, which requires two or more copies of the *RAD51C* gene, would not be expected to occur in PDX models #183 or PH039 in which there is a single copy of the gene. Biallelic inactivation of *BRCA1* achieved by methylation of one allele and somatic loss of the other is an accepted mechanism of functional *BRCA1* loss in *BRCA1* methylated breast and ovarian tumors^32^, and our data now support the relevance of this mechanism for me*RAD51C*.

While all PH039 tumor aliquots lost me*RAD51C* completely following three cycles of rucaparib 450 mg/kg, PDX #183 tumor aliquots had stable methylation patterns. Three of 12 (25%) PDX #183 tumor aliquots presented with a small degree of me*RAD51C* loss following four cycles of rucaparib at 300 mg/kg (Figure 4E; Supplementary Fig. 8A-B; Full summary of MS-HRM screening results found in Supplementary Table 6). Rucaparib 450 mg/kg treatment data also indicated that PDX #183 had relatively stable me*RAD51C* under PARPi pressure compared with either PH039 lineage. Only one PDX #183 tumor aliquot had extensive loss of me*RAD51C* (post-niraparib, Supplementary Fig. 8C-D; Supplementary Table 5). Cisplatin treatment was not as effective as PARPi in driving me*RAD51C* loss in PDX #183 and PH039 lineage B (Supplementary Fig. 8E; Supplementary Table 5). Nonetheless, the niraparib 100 mg/kg and rucaparib 300 mg/kg treatment data established that PH039-A consistently had the least stable me*RAD51C* under PARPi pressure compared with either PDX #183 or PH039-B (Figure 4E; Supplementary Table 6).

Loss of me*RAD51C* correlated with increased/detectable gene expression in rucaparib-treated PH039-A and PH039-B aliquots and niraparib-treated PH039-A (Figure 4F; Supplementary Fig.9A-B). This is consistent with additional findings for this model under niraparib re-treatment pressure ^29^. Me*RAD51C* loss was less frequent in PDX #183, although PDX #183 tumor aliquots with me*RAD51C* loss also had elevated *RAD51C* gene expression, albeit the levels of *RAD51C* expression detected in this PDX were much lower relative to PH039 samples (Supplementary Fig. 9C-D).

Copy number profiles of unmethylated PDX PH039 lineage A and B aliquots were assessed (using SNP arrays) to determine whether unmethylated cells were derived from the same pre-existing clones. It was found that resulting unmethylated PDX aliquots were derived from distinct clones within each lineage, indicating that PARPi resistance was not due to selection of rare pre-existing resistant subclones within the original PH039 HGSC, but was instead acquired independently within each PH039 lineage (Figure 4G).

### Analysis of RAD51C methylation patterns in patient HGSC samples

In order to assess the relevance of the PDX findings for the clinic more broadly, we accessed HGSC patient samples harboring me*RAD51C.* This cohort included three women recruited to the Australian Ovarian Cancer Study (AOCS) as part of the International Cancer Genome Consortium - Ovarian Cancer Project ^33^, one recently diagnosed woman from the Stafford Fox Rare Cancer (SFRC) program (Supplementary Fig. 10), and nine previously reported cases from the University of Washington cohort (as assessed by MS-PCR)^9^. Patients were either chemo-naïve or pre-treated.

### Heterogeneous and homogeneous patterns of RAD51C methylation found in patient HGSC samples

DNA from these 11 patient HGSC samples was tested on both the MS-HRM and targeted me*RAD51C* NGS assays, while the SFRC case was only tested by MS-HRM, revealing both homogeneous and heterogeneous patterns of me*RAD51C* (Table 1; Supplementary Fig. 10-12). Heterogeneous me*RAD51C*, like that of PDX PH039-A, was detected in half of all patient samples (6/12 (50%); Table 1; Figure 5A; Supplementary Fig. 10-12;) with no obvious association with personal/family history, stage, histology, platinum response or previous chemotherapy (Table 1; Supplementary Table 7). Merged NGS amplicon summaries of AOCS and LS patient samples revealed that heterogeneously methylated samples LS473, LS467 and AOCS-120 had reduced methylation at the same CpG sites as PDX PH039-A (CpG −69 and/or −114), while heterogeneously methylated patient sample AOCS-143 had reduced methylation at sites CpG −49 and −93, and LS472 had reduced methylation at sites CpG −43, −49 and −83 (Figure 5A; Supplementary Fig. 12). Importantly, like the me*RAD51C* PDX samples, patient cancers also appeared to have a bimodal distribution of CpG methylation patterns – where epialleles either had > 7/13 CpG sites methylated, or 0 CpG sites methylated (unmethylated epialleles; Figure 5B; Supplementary Fig. 13). Thus, the “% methylated alleles” used to calculate zygosity in Table 1 refers to all methylated alleles detected.

**Figure 5.**
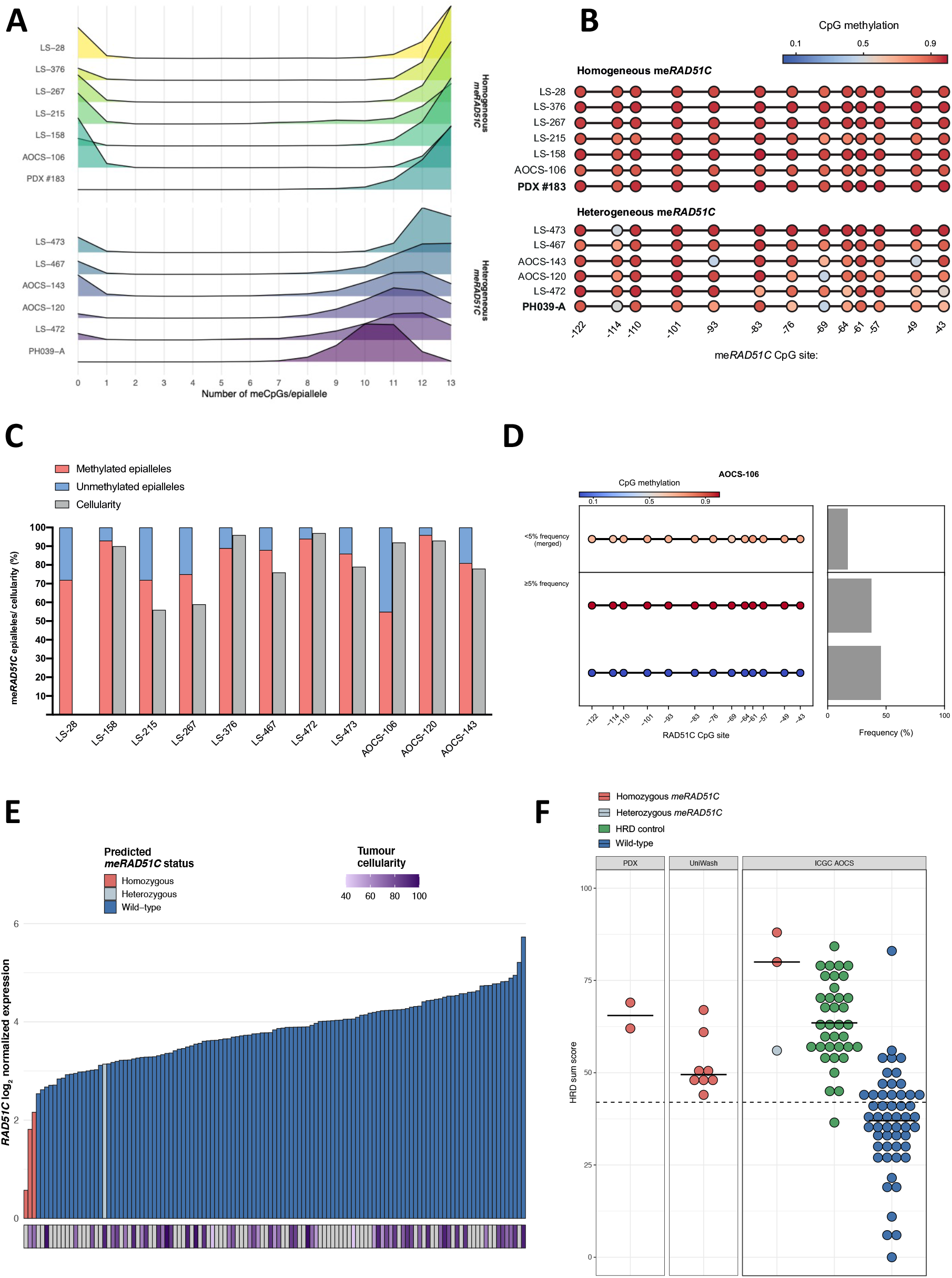
Both homogeneous and heterogeneous me *RAD51C* profiles uncovered in patients. **(A)** Representation of different epialleles based on number of methylated CpG sites (meCpGs) per epiallele. Samples were classified as homogeneous or heterogeneous based on the spread of this distribution for epialleles with >7 meCpGs. **(B)** Average methylation level per CpG site (excluding fully unmethylated epialleles) shows some heterogeneous me*RAD51C* patient samples (like AOCS-143) have reduced methylation of different CpG sites compared to PDX PH039 (PH039-A = PH039 lineage A me*RAD51C* epiallele profile). Positions of CpG sites are relative to the NM_058216 *RAD51C* transcription start site. **(C)** The proportion of methylated/unmethylated epialleles and corresponding tumor purity estimates for each sample (based on qpure SNP array data). LS-28 germline DNA was not available for cellularity estimates. AOCS-143 cellularity was based on a second piece of the same tumor. **(D)** The targeted me*RAD51C* sequencing profile of sample AOCS-106 revealed a homogeneous me*RAD51C* pattern with a high degree of unmethylated epialleles (45% found by sequencing), despite high tumor purity (92.14%). **(E)** RNAseq data from the ICGC study revealed elevated *RAD51C* gene expression in AOCS-106 relative to other me*RAD51C* tumor samples. Bars below (purple scale) indicate tumor purity previously assessed^33^; grey bars indicate tumor purity not available. Order of homozygous samples from left to right: AOCS-120, AOCS-147 (primary sensitive tumor with 100% me*RAD51C* previously assessed by MS-HRM), AOCS-143. RAD51C gene counts – log2(TMM counts + 1)**. (F)** All patient tumors with me*RAD51C* were found to have high HRD sum scores using SNP array data, indicating that these samples were either HRD at time of collection, or prior to. Points above the dashed line are classified as having HRD/history of HRD, bars below dashed line are considered HRR competent with no history of HRD^48, 49, 50^. UniWash (University of Washington) “LS” samples and PDX samples were analyzed with Global Screening Array-24 v2-0 arrays, while ICGC samples were tested as previously described^33^.

Tumor purity and *RAD51C* copy number, estimated from SNP array data using the same DNA samples with me*RAD51C* sequencing available, allowed us to calculate methylation zygosity (Figure 5C; Table 1). All tested samples had high me*RAD51C* (90-100%) when accounting for neoplastic cellularity, except for sample AOCS-106, which had approximately 60% me*RAD51C* (Table 1; Figure 5C). Interestingly, HGSC AOCS-143 also had a pathogenic germline *BRCA1* mutation c.2716A>T (p.Lys906*) detected by sequencing^33^. As this mutation is in the 11q region of exon 10, which is spliced out of the D11q isoform of *BRCA1*, the relative contributions of this mutation and the high me*RAD51C* to HRD in this cancer is unclear^34^.

### Heterozygous RAD51C methylation uncovered in a pre-treated HGSC patient sample with elevated RAD51C gene expression

The adjusted me*RAD51C* measurements for the patient HGSC sample, AOCS-106, suggested a heterozygous me*RAD51C* profile, with one of three *RAD51C* gene copies lacking promoter methylation (60% me*RAD51C* in neoplasm; Figure 5C-D; Table 1). We have previously reported on such a 2/3 (~66%) heterozygous methylation pattern in the *BRCA1*-methylated cell line OVCAR8, which was confirmed to have *BRCA1* expression, HRR capacity and was PARPi-resistant^20^. Thus, this pattern of heterozygous me*RAD51C* was also expected to result in restored *RAD51C* gene expression and HRR proficiency. This was supported by the RNAseq data for this case, where *RAD51C* gene expression was elevated relative to other me*RAD51C* cases (Figure 5E). The higher neoplastic purity of this sample relative to others provides even stronger evidence for restored *RAD51C* gene expression within the carcinoma. This sample was from a patient who had received neo-adjuvant chemotherapy prior to sample collection (1^st^ line carboplatin and paclitaxel). Given that homozygous *BRCA1* methylation has been shown to become heterozygous following chemotherapy in other studies^20^, it is likely that this carcinoma was initially homozygous for me*RAD51C*, subsequently becoming heterozygous (losing methylation of one *RAD51C* copy) under treatment pressure. Although no pre-treatment cancer samples were available for this case to confirm the initial me*RAD51C* status, an HRD score analysis revealed genomic signatures consistent with historic HRD (Figure 5F; Supplementary Fig. 14). Thus, we hypothesize from these data that this HGSC originally had HRD caused by complete me*RAD51C*, which was lost following 1^st^-line chemotherapy.

A second piece of post-chemotherapy sample LS-473 was identified as another potential heterozygous me*RAD51C* sample, with an adjusted neoplastic me*RAD51C* value of 85%, compared to 97% in the first HGSC piece (Supplementary Fig. 13; Supplementary Table 8). Given this cancer was estimated to have 4 copies of the *RAD51C* gene (SNP arrays), loss of methylation in one cellular epiallele (sufficient for competent HRR DNA repair) in all cells would result in tumor me*RAD51C* value of 75%. It is possible that only a portion of cells within the sample had lost methylation of *RAD51C* copy resulting in an adjusted me*RAD51C* value higher than 75% but less than 100%, however we could not assign a heterozygous status to this cancer with confidence. This case highlights the complexity of interpreting me*RAD51C* zygosity in patient HGSC samples.

## DISCUSSION

The impact of *RAD51C* gene methylation on PARPi responses in HGSC has long been unclear^8, 9, 10, 12, 13, 19^. In this study we have demonstrated that me*RAD51C* is a *bona fide* HRR defect which could sensitize HGSC to PARPi treatment, but once lost under treatment pressure, resulted in PARPi-refractory PDX tumors. As seen for me*BRCA1*, a single unmethylated copy of *RAD51C* in HGSC cells was found to be sufficient to restore HRR DNA repair and cause PARPi resistance in PDX. Thus, the presence of fully unmethylated epialleles (those not derived from stroma) should be estimated using tumor purity and *RAD51C* copy number when analyzing me*RAD51C* in HGSC samples.

We described two patterns of me*RAD51C* in HGSC tissues, homogeneous and heterogeneous, and showed both to be associated with gene silencing in PDX and clinical samples, and with PARPi response in PDX. Initially detected in HGSC PDX, heterogeneous me*RAD51C* was confirmed to be present in 50% of me*RAD51C* HGSC patient samples screened. Heterogeneous me*RAD51C* occurs when some CpG sites show no methylation in a given allele and multiple alleles with different methylation patterns co-exist in the region under investigation (reviewed by Mikeska et al., 2010^35^). Although it is possible that some promoter CpG sites are less critical for *RAD51C* silencing than others, more cases with rare me*RAD51C* HGSC would be required to investigate this. However, in our study, only fully unmethylated epialleles were associated with active gene expression, while all silenced epialleles had at least 54% of the region methylated. Thus, any degree of *RAD51C* promoter methylation should be flagged as a potential HRR defect for further validation to ensure that heterogeneously methylated samples with potential for PARPi response are not missed^33^.

Our group has previously demonstrated that loss of methylation of a single methylated *BRCA1* epiallele is sufficient to restore HRR in cells and cause PARPi resistance^20^. In this study, we have confirmed that the same principles apply to *RAD51C* in PDX models #183 and PH039, which by chance both only had one cellular copy of the gene. This single-allele haplo-sufficiency of *RAD51C* was also previously observed by our group for resistance-causing secondary mutations in *RAD51C*^25^. These observations support a model in which heterozygous me*RAD51C* (mix of methylated and unmethylated epialleles within a cell) would also cause gene expression and, thus, PARPi resistance in HGSC. Indeed, our method of me*RAD51C* characterization in patient samples enabled us to detect a pre-treated patient sample with apparent heterozygous me*RAD51C*, where we assessed that two gene copies retained methylation and one had lost methylation. Although matched archival/untreated neoplastic material was not available, all evidence supported a model in which homozygous me*RAD51C* had become heterozygous following chemotherapy. Future studies of me*RAD51C* zygosity in HGSC would benefit from the availability of matched pre- and post-treatment patient HGSC samples.

Importantly, given that loss of me*RAD51C* is associated with restored HRR and PARPi resistance, therapeutic strategies directly causing DNA methylation loss, should be avoided in patients with me*RAD51C* or me*BRCA1*. For example, anti-cancer therapies, such as histone deacetylase (HDAC) and DNA methyltransferase (DNMT) inhibitors may cause permanent me*RAD51C* loss and remove an important therapeutically targetable feature of the patient’s cancer. These compounds are currently being trialed in recurrent ovarian cancers due to their immunogenic effects (e.g. trial NCT02915523)^36^, and their timing should be an important consideration for HGSC patients with known *BRCA1* or *RAD51C* methylation, which collectively account for up to 12% of all HGSC cases.

Through our work on PDX models, we found that the model with the most heterogeneous me*RAD51C* (PDX PH039-A) had the least stable me*RAD51C* under treatment pressure, relative to other models, whilst the most homogeneous me*RAD51C* PDX (#183) had the most stable methylation. This might suggest that heterogeneous me*RAD51C* is more easily lost following platinum/PARPi treatment. However, heterogeneous me*RAD51C* was detected in nearly half of the screened HGSC patient samples, and there did not appear to be an association with initial platinum chemotherapy response status or patient survival. Indeed, patient sample AOCS-120 still contained 100% heterogeneous me*RAD51C* at recurrence following six lines of chemotherapy, demonstrating that other/additional factors are influencing me*RAD51C* stability under platinum and PARPi pressure in HGSC. Identifying these underlying factors could be of great utility and enable design of novel therapeutic strategies to abrogate this form of PARPi resistance. Useful pre-clinical models for this type of study include the two lineages of PDX PH039 where me*RAD51C* loss and PARPi resistance were acquired independently and not via selection of a pre-existing clone. These observations have important implications for the current design of targeted therapy trials for HGSC.

In conclusion, this study identifies both homogeneous and heterogeneous me*RAD51C* as *bona fide* HRR defects that are targetable with PARPi therapy and have potential for sustained response through multiple lines of therapy. Methylation loss can occur in both cases, leading to restored HRR and PARPi resistance and should be considered when making treatment choices with me*RAD51C* HGSC patients.

## METHODS

### Reagents

Niraparib (TesaroBio) was purchased from MedChemExpress, and Rucaparib was provided by Clovis Oncology. Mouse anti-human RAD51C antibody (sc-56214) was purchased from Santa Cruz Biotechnology, goat anti-beta actin (ab8229), Rabbit anti-human RAD51 (ab133534) and mouse anti-human Geminin antibodies (ab104306) from Abcam, Human-specific EpCam-647 (clone VU1D9, #5447) and rabbit anti-human γH2AX antibodies (Phospho-Histone H2A.X Ser139; clone 20E3, #9718) from Cell Signaling Technologies, and EpCam-APC antibody (clone EBA-1, #347200) from BD Biosciences. LIVE/DEAD Fixable Aqua Dead Cell Stain Kit, for 405 nm excitation was purchased from Invitrogen (#L34957). The following antibodies were used for immunohistochemistry: p53 (M700101 1:100; Dako), Ki67 (M7240 1:50; Dako), Cytokeratin (Pan-CK; M3515 1:200; Dako), PAX8 (10336–1-AP 1:20000; Proteintech), and WT1 (ab15249; 1:800; Abcam).

### Study approval

All experiments involving animals were performed according to the animal ethics guidelines and were approved by the Walter and Eliza Hall Institute of Medical Research (WEHI) Animal Ethics Committee. Ovarian carcinoma PDX were generated from patients enrolled in the Australian Ovarian Cancer Study. Informed consent was obtained from all patients, and all experiments were performed according to the human ethics guidelines. Additional ethics approval was obtained from the Human Research Ethics Committees at the Royal Women’s Hospital, the WEHI and QIMR Berghofer (P3456 and P2095).

### Establishment of HGSC Patient Derived Xenografts

HGSC PDX models #183, #1240 and PH039 were established by transplanting fresh fragments (#183, #1240) or viably frozen minced tumor (PH039) subcutaneously into NOD/SCID IL2Rγnull recipient mice (T1, passage 1) as described previously^37^. PDX #183 was from a 65 year-old patient diagnosed with bulky stage III HGSC, and was confirmed to be HGSC by histological staining (Supplementary Fig. 1). PDX was established following surgical de-bulking. This patient was subsequently treated with paclitaxel and carboplatin chemotherapy and bevacizumab in the first-line. The patient responded to first-line treatment, with a treatment-free interval (TFI) of 11 months. Second-line therapy of paclitaxel and carboplatin chemotherapy resulted in a reduced TFI of 7 months. The TFI was shorter again for third-line therapy of carboplatin and gemcitabine, where it was 4 months. This patient died from the disease 3 years and two months from diagnosis, having never received PARPi. PDX #1240 derived from ovary tumor material from primary surgery in a 58 year-old woman diagnosed with stage IIIB HGSC. At last follow-up the patient had experienced a para-aortic (PA) node recurrence less than six months following completion of platinum-based chemotherapy, and was prescribed high dose palliative radiotherapy to the PA region combined with radio-sensitizing cisplatin.

### Treatments of HGSC Patient Derived Xenografts

Recipient mice bearing T2-T6 (passage 2 to passage 6) tumors were randomly assigned to treatments when tumor volume reached 180-300 mm^3^ (PDX #183 and #1240) or 180-250 mm^3^ (faster growing PDX PH039). *In vivo* cisplatin treatments were performed as previously described^37^. The regimen for rucaparib treatment was oral gavage once daily (Monday-Friday) for 3 weeks at 150 mg/kg, 300 mg/kg or 450 mg/kg. The regimen for niraparib treatment was oral gavage once daily (Monday-Friday) for 3 weeks at 100 mg/kg. Tumors were harvested once tumor volume reached 600-700 mm^3^ or when mice reached ethical endpoint. Rucaparib re-treatment experiments in PDX #183 involved standard 300 mg/kg treatment of rucaparib followed by an additional two weeks of 300 mg/kg rucaparib treatment (oral gavage once daily, Monday-Friday) once tumor volume reached 500 mm^3^. Tumors were then transplanted into a second recipient mouse where the treatment was repeated for a total of four treatment cycles (two per mouse). Rucaparib re-treatment experiments in PDX PH039 involved standard 450 mg/kg treatment of rucaparib followed by an additional two weeks of 450 mg/kg rucaparib treatment (oral gavage once daily, Monday-Friday) once tumor volume reached 300 mm^3^. Tumors were then transplanted into a second recipient mouse where the treatment was repeated for a total of four treatment cycles (two per mouse). Nadir, time to progression (TTP or PD), time to harvest (TTH), and treatment responses are as defined previously^20^.

### Immunohistochemistry

Automated immunohistochemical (IHC) staining was performed on the DAKO Omnis (Agilent Pathology Solutions) samples to confirm HGSC characteristics of PDX.

### Immunoblotting

PDX tumor protein lysates were prepared using 2D lysis buffer containing protease inhibitor PMSF (7M urea, 2M Thiourea, 4% CHAPS in 0.1M Tris buffer with 4.6% 100 mM PMSF). Equal protein loads of 8ug were resolved on precast 8% to 16% Bis-Tris gels (Bio-Rad) under reducing conditions. Protein was transferred to PVDF membranes using Bio-Rad Criterion wet transfer method (Bio-Rad), then probed with anti-RAD51C (Santa Cruz, sc-56214) followed by peroxidase-labeled secondary antibody (SouthernBiotech, 1010-05). Membranes were then stripped and probed with anti-beta actin (Abcam), followed by peroxidase-labeled secondary antibody (Santa Cruz, sc-2020) for loading control. Blots were visualized by enhanced chemiluminescence (Amersham ECL Prime, GE Healthcare Life Sciences) according to manufacturer’s instructions.

### DNA methylation by MS-PCR

DNA was extracted from tumor samples using the QIAamp DNA Mini Kit (Qiagen) according to manufacturer’s instructions. 250 ng of DNA was bisulfite converted (EZ Methylation Direct kit, Zymo Research, Irvine, CA) and evaluated with methylation sensitive PCR for *RAD51C* as previously described ^12^.

### DNA methylation by MS-HRM

Patterns of me*RAD51C* were assessed by MS-HRM as previously described^38^ on the CFX (Bio-Rad) or MIC (bio molecular systems) thermocycler platforms, using primers targeting the *RAD51C* promoter (Methods Table 1) at genomic region chr 17:56,769,849-56,769,990 (hg19)^38^.

### Targeted RAD51C bisulfite amplicon sequencing

The primers used in *RAD51C*-targeted MS-HRM were modified by the addition of Illumina sequencing adaptors to create an NGS-based assay that would allow interrogation of individual CpG sites and amplicons (alleles) within me*RAD51C* PDX samples. Bisulfite-converted DNA (bisDNA) samples were not quantitated prior to PCR, but was instead quantitated and normalized prior to sequencing.

For the inner PCR reaction, 4 μl of bisDNA was added to 2 μl 10x PCR buffer (Qiagen, Cat# 203203), 2 μl of 10 μM primer mix (Methods Table 1), 0.4 μl of 10 μM dNTP mix, 11.5 μl of molecular grade H2O and 0.1 μl HotStarTaq DNA Polymerase (Qiagen, Cat# 203203). Reactions were incubated at 95°C for 15 minutes, followed by 15 cycles of 95°C for 30 seconds, 60°C for 40 seconds, 72°C for 40 seconds. Reactions then incubated at 72°C for 10 minutes, and stored at 4°C. In the outer PCR, unique combinations of Nextera XT Index Kit v2 primers (Illumina) were added for sample multiplexing. Briefly, 2 μl of inner PCR product was added to 2 μl 10x PCR buffer (Qiagen, Cat# 203203), 0.4 μl of 10 μM dNTP mix, 13.5 μl of molecular grade H_2_O, 0.1 μl HotStarTaq DNA Polymerase (Qiagen, Cat# 203203) and 1 μl of each Nextera XT Index Kit v2 set B or set C barcoding primer (Illumina, Cat# FC-131-2002 or FC-131-2003). These reactions were then incubated at 95°C for 15 minutes, followed by 20 cycles of 94°C for 30 seconds, 60°C for 40 seconds, 72°C for 40 seconds. Reactions were then incubated at 72°C for 10 minutes, and stored at 4°C. Resulting libraries were cleaned-up using Agencourt AMPure XP (Beckman Coulter, Cat#10136224) beads using a ratio of 0.9:1. The D1000 ScreenTape System (Agilent, Cat#5067-5582 and 5067-5583) was used to assess the size profile of NexteraXT libraries according to manufacturer’s protocol. NGS libraries were quantitated using the Qubit dsDNA high sensitivity (HS) assay, normalized to 2 nM and pooled. These pools were then sequenced in the WEHI Genomics Core Laboratory on the Illumina MiSeq using a MiSeq Nano Reagent Kit v2 (300 cycle; Illumina; Cat# MS-102-2002) according to manufacturer’s protocol. Libraries were sequenced to a minimum depth of 5,000. Demultiplexed reads were merged and grouped using AmpliVar genotyping mode^39^, then processed using a custom script, available on github (https://github.com/okon/MethAmplicons). Reads with deletions, insertions or substitution variants in the CpG positions were filtered out. Epialleles below 5% frequency were grouped together for individual epiallele plots as they were more likely to contain sequencing or bisulfite conversion errors.

### BROCA sequencing

BROCA-Cancer Risk panel sequencing was performed and analyzed as previously described^26^ using version BROCA-HR v7 of the panel (http://tests.labmed.washington.edu/BROCA).

### RNA expression analysis in PDX tumors

RNA was isolated from snap-frozen tissue using the Direct-zol RNA MiniPrep kit (Zymo Research, Cat# R2050) and quality was assessed using the RNA ScreenTape System (Agilent) according to manufacturer’s protocol. The qRT-PCR was performed in triplicate using 100 ng of RNA and the TaqMan RNA-to-CT 1-Step Kit (Applied Biosystems, Carlsbad, CA) per manufacturer’s instructions. Using GAPDH (4352665, Life Technologies) and RAD51C (Hs04194939_s, Applied Biosystems, Carlsbad, CA) probe sets, PCR was performed on a CFX384 Real Time System (C10000 Touch Thermal Cycler, BioRad, Hercules, CA) using a program consisting of 48°C for 15 min, 95°C for 10 min, then 40 cycles of 95°C for 15 sec and 60°C for 1 min. Data were analyzed using the following equations: ΔC_t_=C_t_(sample)-C_t_(endogenous control); and fold-change = 2^−ΔCt^. Relative *RAD51C* fold-change expression values were normalized to PEO1 cell line control for comparison. RNA sequencing library preparation and data analysis were performed as previously described^20^.

### RAD51 foci formation in PDX tumors

PDX tumors were either harvested (untreated) or irradiated (*in vivo* at 5Gy) and harvested 4 hours post-irradiation at a 400-700 mm^3^ volume. Tumors were dissociated by mincing followed by a 45 minute incubation at 37°C in digestion media (DMEM/F-12 with GlutaMAX (Gibco), 10 mg/ml Pronase (Sigma-Aldrich) and 1mg/ml DNAse (Sigma-Aldrich)). After digestion, cells were passed through a 100μm cell strainer to create single-cell suspensions. Cells were then stained with LIVE/DEAD Fixable Aqua Dead Cell Stain according to manufacturer’s instructions, and blocked in 2.4G2 supernatant diluted 1:2 in 3% FCS in DPBS, or 7% FCS in DPBS, for a 15 minutes on ice. Cells were stained with human-specific EpCam-647 antibody or EpCam-APC for a minimum dilution of 1:100 or 1:20 antibody respectively. Cells were then fixed with 200-1000μl of 4% Paraformaldehyde for 5 minutes. Cells were sorted for the EpCam-647/APC positive (epithelial cells) and LIVE/DEAD-aqua negative (alive following digestion) population.

Resulting cells were ultra-concentrated in 10 μl of DPBS and plated onto wells of a CellCarrier-96 Ultra Microplates (PerkinElmer, Cat# 6055302). Plates were then centrifuged at 1800g for 10 minutes to allow the cells stick to the surface of the wells. They were then immediately permeabilized using 0.2% Triton-X-100, incubated for 10 minutes at room temperature. Cells were washed (3 times with 150 μl DPBS per well), and blocked in IFF buffer (DPBS with 2% FCS and 1% BSA) for 1 hour (room temperature). Cells were then washed and incubated with anti-Geminin antibody diluted 1:100 in IFF buffer for 1 hour (room temperature). Cells were washed again and incubated overnight at 4°C with either anti-RAD51 or anti-γH2AX antibody, diluted 1:250 and 1:200 in IFF buffer respectively. The following day, cells were washed and incubated for 30 minutes at room temperature with a solution of Goat anti-Rabbit IgG (H+L) Cross-Adsorbed Secondary Antibody, Alexa Fluor 488 (Invitrogen, Cat# A-11008) and Goat anti-Mouse IgG (H+L) Highly Cross-Adsorbed Secondary Antibody, Alexa Fluor 546 (Invitrogen, Cat# A-11030) – both diluted 1:800 in IFF buffer, combined with 1 drop of Hoechst 33342 Ready Flow™ Reagent (Invitrogen, Cat# R37165) per ml of solution. Cells were then washed once in 150 μl of DPBS and left in 150 μl of DPBS until imaging.

### DNA repair foci imaging and analysis

Cells stained for DNA repair foci on CellCarrier-96 Ultra Microplates (PerkinElmer, Cat# 6055302) were imaged on the Opera Phenix™ High Content Screening System (PerkinElmer). The imaging settings were as follows: optical mode was set to confocal with an imaging plane of 1.5 μm using a 40x water immersion. There were 197 fields of view captured per well. The channels Alexa 488, Alexa 568 and HOECHST 33342 were used to measure RAD51/γH2AX, geminin and nuclear stains respectively. Files were then exported automatically to the WEHI Columbus server (PerkinElmer), where aa custom foci analysis pipeline was run using the following settings: Find nuclei was set to method C with a >30 μm^2^ area threshold and standard morphology properties. Nuclei were then filtered based on an area of 50-150μm^2^ and a roundness of >0.9 to make the “assay nuclei” group. This group was then further filtered based on a geminin staining intensity mean (Alexa 568) of >800-1500. The “find spots” protocol was then applied, using method C to filter for spots with a mean intensity of >900-2500 within the geminin positive group. The results were plotted using the PRISM7 (GraphPad) software.

### 450K methylation array analysis

Methylation data from and publicly available 450K array data (GEO accession number GSE65820) for AOCS patient samples were analyzed using the Miss Methyl package in R ^40^. The CpG sites used to assess me*RAD51C* were based on the *RAD51C* promoter region covered by the MS-HRM assay: (cg05214530 is NM_058216.2:c.-114/ chr17:g.56769891 and cg27221688 is NM_058216.2:c.-101/ chr17:g.56769904).

### Patient Samples

Four samples from the AOCS cohort were identified to have me*RAD51C* using publicly available methylation array data (three samples were previously reported as methylated)^33^. We were able to source tumor and germline DNA for three of the four cases (AOCS-106, AOCS-143 and AOCS-120). We also sourced tumor and germline DNA from an additional nine previously published me*RAD51C* samples (LS samples) ^9^. DNA from these samples was tested by MS-HRM and targeted me*RAD51C* bisulfite sequencing (for quantitative me*RAD51C* assessment), as well as SNP arrays to determine tumor purity and *RAD51C* gene copy number.

### Single nucleotide polymorphism arrays

Single nucleotide polymorphism (SNP) arrays were performed on the Global Screening Array-24 v2-0 (Illumina) and scanned on an iScan (Illumina) as previously described^33^. Data were processed using the Genotyping module (v.1.9.4) in GenomeStudio v.2011.1 (Illumina) to calculate B-allele frequencies (BAF) and logR ratios. Copy number changes were estimated using GAP^41^, after low-quality probes assessed in the matched normal sample with GenCall (GC) score of <0.7 were removed (where matched normal sample was available). Tumor cellularity was estimated using qpure^42^.

### Adjusted tumor meRAD51C calculation

The following formula was used to calculate the adjusted (tumor) me*RAD51C*:

Adjusted me*RAD51C* = Observed me*RAD51C* % × Total *RAD51C* copy number ÷ Tumor *RAD51C* copy number × Tumour cellularity

Where Total *RAD51C* copies = Tumour cellularity % × Tumor *RAD51C* copy number + (100 – Tumour cellularity %) × 2

### RNA sequencing analysis of public data

Publicly available RNA sequencing data for AOCS patient samples^33^ was analyzed using the standard pipeline. Briefly, adapter sequences were trimmed using Cutadapt (version 1.11^43^) and aligned using STAR (version 2.5.2a^44^) to the GRCh37 assembly with the gene, transcript, and exon features of Ensembl (release 75) gene model. Quality control metrics were computed using RNA-SeQC (version 1.1.8^45^) and expression was estimated using RSEM (version 1.2.30^46^). Data was corrected for library size using counts per million (CPM) and was corrected for differences in RNA composition using trimmed mean of M values (TMM^47^).

### HRD score analysis

HRR-deficiency (HRD) scores were assessed on SNP array data using scarHRD package on the allele-specific copy number calls. The HRD sum score cut-off of ≥ 42 was used to define HRD, based on the previous reports in breast and ovarian cancer^48, 49^.

## ACKNOWLEDGMENTS

We thank Dr. Paul Haluska and Mariam AlHilli (Mayo Clinic, USA) for the cryopreserved PDX material used to re-establish PDX PH039 within our laboratory, and for original PH039 characterization. We also thank Silvia Stoev, Rachel Hancock, Kathy Barber, Scott Wood and Conrad Leonard for technical assistance. We thank Clovis Oncology for providing rucaparib for *in vivo* experiments. This work was supported by fellowships and grants from the National Health and Medical Research Council (NHMRC Australia; Project grant 1062702 (CLS)); the Stafford Fox Medical Research Foundation (CLS); Cancer Council Victoria (Sir Edward Dunlop Fellowship in Cancer Research to CLS); the Victorian Cancer Agency (Clinical Fellowships to CLS CRF10–20, CRF16014; GD ECRF19003) National Institute of Health (2P50CA083636) (to EMS) and the Wendy Feuer Ovarian Cancer Research Fund (to EMS); the Bev Gray Ovarian Cancer Scholarship (PhD top-up scholarship) and Research Training Program Scholarship (PhD Scholarship) to KN. The Olivia Newton-John Cancer Research Institute acknowledges the support of the Victorian Government Operational and Infrastructure Support Program. This work was also supported in part by grants from the NIH (P50 CA136393 to S.H.K. and S.J.W.; F30 CA213737 to C.D.M.; T32 to GM072474 to R.M.H.), Stand Up to Cancer (E.M.S., S.J.W. and S.H.K.) and fellowship support from the Mayo Foundation for Education and Research (R.M.H., C.D.M.). This work was made possible through the Australian Cancer Research Foundation, the Victorian State Government Operational Infrastructure Support and Australian Government NHMRC IRIISS. All authors, the WEHI Stafford Fox Rare Cancer Program and the AOCS would like to thank all of the women who participated in these research programs. The AOCS would also like to acknowledge the contribution of the study nurses, research assistants, and all clinical and scientific collaborators to the study. The complete AOCS Study Group can be found at www.aocstudy.org. The Australian Ovarian Cancer Study gratefully acknowledges additional support from Ovarian Cancer Australia and the Peter MacCallum Foundation. The Australian Ovarian Cancer Study Group was supported by the U.S. Army Medical Research and Materiel Command under DAMD17-01-1-0729, The Cancer Council Victoria, Queensland Cancer Fund, The Cancer Council New South Wales, The Cancer Council South Australia, The Cancer Council Tasmania and The Cancer Foundation of Western Australia (Multi-State Applications 191, 211 and 182) and the National Health and Medical Research Council of Australia (NHMRC; ID199600; ID400413 and ID400281).

## CONSORTIA

*Australian Ovarian Cancer Study (AOCS)*

Chenevix-Trench G., Green A., Webb P., Gertig D., Fereday S., Moore S., Hung J., Harrap K., Sadkowsky T., Pandeya N., Malt M., Mellon A., Robertson R., Vanden Bergh T., Jones M., Mackenzie P., Maidens J., Nattress K., Chiew Y. E., Stenlake A., Sullivan H., Alexander B., Ashover P., Brown S., Corrish T., Green L., Jackman L., Ferguson K., Martin K., Martyn A., Ranieri B., White J., Jayde V., Mamers P., Bowes L., Galletta L., Giles D., Hendley J., Schmidt T., Shirley H., Ball C., Young C., Viduka S., Tran H., Bilic S., Glavinas L., Brooks J., Stuart-Harris R., Kirsten F., Rutovitz J., Clingan P., Glasgow A., Proietto A., Braye S., Otton G., Shannon J., Bonaventura T., Stewart J., Begbie S., Friedlander M., Bell D., Baron-Hay S., Ferrier A., Gard G., Nevell D., Pavlakis N., Valmadre S., Young B., Camaris C., Crouch R., Edwards L., Hacker N., Marsden D., Robertson G., Beale P., Beith J., Carter J., Dalrymple C., Houghton R., Russell P., Links M., Grygiel J., Hill J., Brand A., Byth K., Jaworski R., Harnett P., Sharma R., Wain G., Ward B., Papadimos D., Crandon A., Cummings M., Horwood K., Obermair A., Perrin L., Wyld D., Nicklin J., Davy M., Oehler M. K., Hall C., Dodd T., Healy T., Pittman K., Henderson D., Miller J., Pierdes J., Blomfield P., Challis D., McIntosh R., Parker A., Brown B., Rome R., Allen D., Grant P., Hyde S., Laurie R., Robbie M., Healy D., Jobling T., Manolitsas T., McNealage J., Rogers P., Susil B., Sumithran E., Simpson I., Phillips K., Rischin D., Fox S., Johnson D., Lade S., Loughrey M., O’Callaghan N., Murray W., Waring P., Billson V., Pyman J., Neesham D., Quinn M., Underhill C., Bell R., Ng L. F., Blum R., Ganju V., Hammond I., Leung Y., McCartney A., Buck M., Haviv I., Purdie D., Whiteman D. & Zeps N.

## AUTHOR CONTIRBUTIONS

C.L.S., M.J.W., A.D., and S.H.K. designed the study, developed the methodology, analyzed the data, wrote the manuscript, and supervised the study.

K.N. and O.K. designed the study, developed the methodology, performed the experiments, analyzed the data, wrote the manuscript, and supervised the study.

R.H. and C.mcG. assisted in study design, analyzed the data, assisted in cross-validation and reviewed the manuscript.

C.J.V. assisted in study design, analyzed the data and reviewed the manuscript.

G.H. and E.L. developed models, performed the experiments and reviewed the manuscript.

G.D., N. B., and K.S.A. performed experiments and reviewed the manuscript.

M.R., A.M., Z.Q.C., M.R.E.G., M.I.H., and K.Nones performed experiments, analyzed data and reviewed the manuscript.

N.T. acquired the data, provided administrative support, and reviewed the manuscript.

A.deF., D.D.B, E.M.S., and S.J.W. acquired the data, supervised the study, and reviewed the manuscript.

N.W. supervised the study, provided administrative support and reviewed the manuscript. and AOCS acquired the data and reviewed the manuscript.

## COMPETING INTERESTS

CLS declares Advisory Boards for AstraZeneca, Clovis Oncology, Roche, Eisai Inc, Sierra Oncology, Takeda, MSD and Grant/Research support from Clovis Oncology, Eisai Inc, Sierra Oncology, Roche and Beigene. A.deF. has received research grant support from AstraZeneca. DK has received funding from Novartis, Bristol-Meyer Squibb and Merck Sharp Dohme for educational presentations and advisory boards. The remaining authors declare no competing interests.

## REFERENCES

1. Moore K, et al. Maintenance Olaparib in Patients with Newly Diagnosed Advanced Ovarian Cancer. N Engl J Med 379, 2495–2505 (2018).

2. González-Martín A, et al. Niraparib in Patients with Newly Diagnosed Advanced Ovarian Cancer. New England Journal of Medicine 381, 2391–2402 (2019).

3. Vaughan S, et al. Rethinking ovarian cancer: recommendations for improving outcomes. Nat Rev Cancer 11, 719–725 (2011).

4. Bowtell DD, et al. Rethinking ovarian cancer II: reducing mortality from high-grade serous ovarian cancer. Nat Rev Cancer 15, 668–679 (2015).

5. Chiang JW, Karlan BY, Baldwin RL. BRCA1 promoter methylation predicts adverse ovarian cancer prognosis. Gynecologic oncology 101, 403–410 (2006).

6. Stefansson OA, Villanueva A, Vidal A, Martí L, Esteller M. BRCA1 epigenetic inactivation predicts sensitivity to platinum-based chemotherapy in breast and ovarian cancer. Epigenetics 7, 1225–1229 (2012).

7. Cunningham JM, et al. Clinical characteristics of ovarian cancer classified by BRCA1, BRCA2, and RAD51C status. Scientific reports 4, 4026 (2014).

8. Ruscito I, et al. BRCA1 gene promoter methylation status in high-grade serous ovarian cancer patients–a study of the tumour Bank ovarian cancer (TOC) and ovarian cancer diagnosis consortium (OVCAD). European journal of cancer 50, 2090–2098 (2014).

9. Bernards SS, et al. Clinical characteristics and outcomes of patients with BRCA1 or RAD51C methylated versus mutated ovarian carcinoma. Gynecologic Oncology 148, 281–285 (2018).

10. Swisher EM, et al. Methylation and protein expression of DNA repair genes: association with chemotherapy exposure and survival in sporadic ovarian and peritoneal carcinomas. Mol Cancer 8, 48 (2009).

11. Cancer Genome Atlas Research N, et al. Integrated genomic characterization of endometrial carcinoma. Nature 497, 67–73 (2013).

12. Swisher EM, et al. Rucaparib in relapsed, platinum-sensitive high-grade ovarian carcinoma (ARIEL2 Part 1): an international, multicentre, open-label, phase 2 trial. Lancet Oncol 18, 75–87 (2017).

13. Lheureux S, et al. Long-Term Responders on Olaparib Maintenance in High-Grade Serous Ovarian Cancer: Clinical and Molecular Characterization. Clin Cancer Res 23, 4086–4094 (2017).

14. Drew Y, et al. Therapeutic potential of poly(ADP-ribose) polymerase inhibitor AG014699 in human cancers with mutated or methylated BRCA1 or BRCA2. J Natl Cancer Inst 103, 334–346 (2011).

15. Min A, et al. RAD51C-deficient cancer cells are highly sensitive to the PARP inhibitor olaparib. Molecular cancer therapeutics 12, 865–877 (2013).

16. Veeck J, et al. BRCA1 CpG island hypermethylation predicts sensitivity to poly(adenosine diphosphate)-ribose polymerase inhibitors. J Clin Oncol 28, e563–564; author reply e565-566 (2010).

17. George J, et al. Nonequivalent gene expression and copy number alterations in high-grade serous ovarian cancers with BRCA1 and BRCA2 mutations. Clinical Cancer Research 19, 3474–3484 (2013).

18. Polak P, et al. A mutational signature reveals alterations underlying deficient homologous recombination repair in breast cancer. Nat Genet 49, 1476–1486 (2017).

19. TCGA CGARN. Integrated genomic analyses of ovarian carcinoma. Nature 474, 609 (2011).

20. Kondrashova O, et al. Methylation of all BRCA1 copies predicts response to the PARP inhibitor rucaparib in ovarian carcinoma. Nat Commun 9, 3970 (2018).

21. Maykel J, Liu JH, Li H, Shultz LD, Greiner DL, Houghton J. NOD-scidIl2rgtm1Wjland NOD-Rag1nullIl2rgtm1Wjl: A Model for Stromal Cell–Tumor Cell Interaction for Human Colon Cancer. Digestive Diseases and Sciences 59, 1169–1179 (2014).

22. Meindl A, et al. Germline mutations in breast and ovarian cancer pedigrees establish RAD51C as a human cancer susceptibility gene. Nat Genet 42, 410–414 (2010).

23. Pelttari LM, et al. RAD51C is a susceptibility gene for ovarian cancer. Human Molecular Genetics 20, 3278–3288 (2011).

24. Loveday C, et al. Germline RAD51C mutations confer susceptibility to ovarian cancer. Nat Genet 44, 475–476; author reply 476 (2012).

25. Kondrashova O, et al. Secondary Somatic Mutations Restoring RAD51C and RAD51D Associated with Acquired Resistance to the PARP Inhibitor Rucaparib in High-Grade Ovarian Carcinoma. Cancer Discov 7, 984–998 (2017).

26. Walsh T, et al. Mutations in 12 genes for inherited ovarian, fallopian tube, and peritoneal carcinoma identified by massively parallel sequencing. Proc Natl Acad Sci U S A 108, 18032–18037 (2011).

27. Pettitt SJ, et al. Genome-wide and high-density CRISPR-Cas9 screens identify point mutations in PARP1 causing PARP inhibitor resistance. Nat Commun 9, 1849 (2018).

28. AlHilli MM, et al. In vivo anti-tumor activity of the PARP inhibitor niraparib in homologous recombination deficient and proficient ovarian carcinoma. Gynecol Oncol 143, 379–388 (2016).

29. Hurley RM, et al. Characterization of a RAD51C-Silenced High Grade Serous Ovarian Cancer Model During PARP Inhibitor Resistance Development. bioRxiv, 2020.2012.2025.419978 (2020).

30. Candiloro ILM, Mikeska T, Hokland P, Dobrovic A. Rapid analysis of heterogeneously methylated DNA using digital methylation-sensitive high resolution melting: application to the CDKN2B (p15) gene. Epigenetics & chromatin 1, 7–7 (2008).

31. Cameron EE, Baylin SB, Herman JG. p15(INK4B) CpG island methylation in primary acute leukemia is heterogeneous and suggests density as a critical factor for transcriptional silencing. Blood 94, 2445–2451 (1999).

32. Esteller M, et al. Promoter hypermethylation and BRCA1 inactivation in sporadic breast and ovarian tumors. J Natl Cancer Inst 92, 564–569 (2000).

33. Patch AM, et al. Whole-genome characterization of chemoresistant ovarian cancer. Nature 521, 489–494 (2015).

34. Wang Y, et al. The BRCA1-Delta11q Alternative Splice Isoform Bypasses Germline Mutations and Promotes Therapeutic Resistance to PARP Inhibition and Cisplatin. Cancer Res 76, 2778–2790 (2016).

35. Mikeska T, Candiloro IL, Dobrovic A. The implications of heterogeneous DNA methylation for the accurate quantification of methylation. Epigenomics 2, 561–573 (2010).

36. Moufarrij S, et al. Epigenetic therapy for ovarian cancer: promise and progress. Clin Epigenetics 11, 7–7 (2019).

37. Topp MD, et al. Molecular correlates of platinum response in human high-grade serous ovarian cancer patient-derived xenografts. Mol Oncol 8, 656–668 (2014).

38. Snell C, Krypuy M, Wong EM, kConFab i, Loughrey MB, Dobrovic A. BRCA1 promoter methylation in peripheral blood DNA of mutation negative familial breast cancer patients with a BRCA1 tumour phenotype. Breast cancer research : BCR 10, R12–R12 (2008).

39. Hsu AL, et al. AmpliVar: mutation detection in high-throughput sequence from amplicon-based libraries. Hum Mutat 36, 411–418 (2015).

40. Phipson B, Maksimovic J, Oshlack A. missMethyl: an R package for analyzing data from Illumina’s HumanMethylation450 platform. Bioinformatics 32, 286–288 (2016).

41. Popova T, Manié E, Stoppa-Lyonnet D, Rigaill G, Barillot E, Stern MH. Genome Alteration Print (GAP): a tool to visualize and mine complex cancer genomic profiles obtained by SNP arrays. Genome Biology 10, R128 (2009).

42. Song S, et al. qpure: A Tool to Estimate Tumor Cellularity from Genome-Wide Single-Nucleotide Polymorphism Profiles. PLOS ONE 7, e45835 (2012).

43. Martin M. Cutadapt removes adapter sequences from high-throughput sequencing reads. EMBnetjournal; Vol 17, No 1: Next Generation Sequencing Data AnalysisDO - 1014806/ej171200, (2011).

44. Dobin A, et al. STAR: ultrafast universal RNA-seq aligner. Bioinformatics 29, 15–21 (2012).

45. DeLuca DS, et al. RNA-SeQC: RNA-seq metrics for quality control and process optimization. Bioinformatics 28, 1530–1532 (2012).

46. Li B, Dewey CN. RSEM: accurate transcript quantification from RNA-Seq data with or without a reference genome. BMC Bioinformatics 12, 323 (2011).

47. Robinson MD, Oshlack A. A scaling normalization method for differential expression analysis of RNA-seq data. Genome Biology 11, R25 (2010).

48. Mills GB, et al. Homologous recombination deficiency score shows superior association with outcome compared with its individual score components in platinum-treated serous ovarian cancer. Gynecologic Oncology 141, 2–3 (2016).

49. Telli ML, et al. Homologous Recombination Deficiency (HRD) Score Predicts Response to Platinum-Containing Neoadjuvant Chemotherapy in Patients with Triple-Negative Breast Cancer. Clin Cancer Res 22, 3764–3773 (2016).

50. Sztupinszki Z, et al. Migrating the SNP array-based homologous recombination deficiency measures to next generation sequencing data of breast cancer. npj Breast Cancer 4, 16 (2018).

