## Supplementary Tables and Figures for "Acquired *RAD51C* promoter methylation loss causes PARP inhibitor resistance in high grade serous ovarian carcinoma"

Nesic *et al.*

**Supplementary Table 1. PDX #183 and #1240 in vivo treatment response summary.** TTH – time to harvest; TTP – time to progression; SD – stable disease; CR – complete response.

| | |
|---|---|
| **PDX #183 vehicle values** | |
| **Median TTH (days)** | 50 |
| **Average TTP (days)** | 8 |
| **PDX #183 cisplatin response** | |
| **Response** | Sensitive (CR) |
| **Median TTH (days)** | 183 |
| **Average TTP (days)** | 152 |
| ***p*-value** | <0.0001 |
| **PDX #183 rucapraib 300 mg/kg response** | |
| **Response** | Responsive (SD) |
| **Median TTH (days)** | 74 |
| **Average TTP (days)** | 36 |
| ***p*-value** | 0.01 |
| **PDX #183 rucapraib 450 mg/kg response** | |
| **Response** | Responsive (SD) |
| **Median TTH (days)** | 95 |
| **Average TTP (days)** | 47 |
| ***p*-value** | <0.0001 |
| **PDX #183 niraparib 100 mg/kg response** | |
| **Response** | Responsive (SD) |
| **Median TTH (days)** | 95 |
| **Average TTP (days)** | 50 |
| ***p*-value** | <0.0001 |
| **PDX #1240 vehicle values** | |
| **Median TTH (days)** | 25 |
| **Average TTP (days)** | 4 |
| **PDX #1240 cisplatin response** | |
| **Response** | Sensitive (CR) |
| **Median TTH (days)** | 130 |
| **Average TTP (days)** | >130 |
| ***p*-value** | <0.0001 |
| **PDX #1240 rucapraib 300 mg/kg response** | |
| **Response** | Responsive (SD) |
| **Median TTH (days)** | 109 |
| **Average TTP (days)** | 81 |
| ***p*-value** | 0.01 |

**Supplementary Table 2. BROCA sequencing results for case #183 and #1240 patient and PDX samples.** Both #183 patient and PDX samples carry a deleterious TP53 mutation and loss of one copy of PTEN. The heterozygous PARP1 mutation detected in the PDX affects the regulatory domain of the protein. In silico prediction in mutation taster (Schwarz et al., 2014) indicated that this mutation is pathogenic, but it was not reported as a PARPi-resistance mutation in published CRISPR screens (Pettitt et al., 2018). Thus the effects of this PARP1 mutation on PARPi activity remain unclear. ExAC = The Exome Aggregation Consortium; 1000g = 1000 genomes.

| Variant-type (short variant, CNV, rearrangement) | Gene | Transcript | c.DNA change | Protein change | Genomic position (hg19) | Coverage (reads) | % Variant Allele | Copy Number Estimate | CNV exons | CNV type | CNV reads | ExAC / 1000 genomes frequencies | In silico prediction |
|---|---|---|---|---|---|---|---|---|---|---|---|---|---|
| short-variant | *TP53* | NM_000546 | c.661G>T | p.E221X | chr17:7578188 | 182 | 64% | | | | | Absent in both | Pathogenic |
| CNV | *PTEN* | NM_000314 | | | chr10:89592934-89718404 | | | 1 | 1-7 | LOSS | 30\|16 | | |
| short-variant | *TP53* | NM_000546 | c.661G>T | p.E221X | chr17:7578188 | 105 | 99% | | | | | Absent in both | Pathogenic |
| short-variant | *PARP1* | NM_001618 | c.2008C>A | p.Q670K | chr1:226561989 | 254 | 51% | | | | | Absent in both | Pathogenic |
| CNV | *PTEN* | NM_000314 | | | chr10:89592934-89718404 | | | 1 | 1-7 | LOSS | 52\|19 | | |
| short-variant | *ATM* | NM_000051 | c.2638+4A>G | | chr11:108138073 | 489 | 0.65 | | | | | Absent in both | Pathogenic |
| short-variant | *XRCC4* | NM_022550 | c.970A>C | p.N324H | chr5:82649026 | 636 | 0.61 | | | | | ExAC: 0.0002<br>1000g: 0.000199681 | Pathogenic |
| short-variant | *CHEK1* | NM_001114122 | c.829C>T | p.R277X | chr11:125513701 | 482 | 0.33 | | | | | ExAC: 0.00003793<br>Absent in 1000g | Assumed pathogenic (stopgain) |
| CNV | *FANCL* | NM_018062 | c.156_1220del | p.R52fs | chr2:58386808-58458714 | | | 1 | 3-14 | DEL | 446\|60 | | |
| short-variant | *TP53* | NM_000546 | c.920-1G>A | | chr17:7576927 | 494 | 0.76 | | | | | Absent in both | Pathogenic |

**Supplementary Table 3. PDX PH039-A and PH039-B in vivo treatment response summary.**

TTH – time to harvest; TTP – time to progression; SD – stable disease; CR – complete response.

| **PDX PH039-A vehicle values** | | **PDX PH039-B vehicle values** | |
|---|---|---|---|
| **Median TTH (days)** | 8 | **Median TTH (days)** | 15 |
| **Average TTP (days)** | 4 | **Average TTP (days)** | 4 |
| ***p*-value comparing lineages** | | 0.0148 | |
| **PDX PH039-A cisplatin response** | | **PDX PH039-B cisplatin response** | |
| **Response** | Sensitive (CR) | **Response** | Sensitive (CR) |
| **Median TTH (days)** | >120 | **Median TTH (days)** | 109 |
| **Average TTP (days)** | 134 | **Average TTP (days)** | 106 |
| ***p*-value** | 0.0008 | ***p*-value** | 0.001 |
| ***p*-value comparing lineages** | | 0.1269 | |
| **PDX PH039-A rucapraib 300 mg/kg response** | | **PDX PH039-B rucapraib 300 mg/kg response** | |
| **Response** | Responsive (SD) | **Response** | Responsive (PR) |
| **Median TTH (days)** | 53 | **Median TTH (days)** | 67 |
| **Average TTP (days)** | 43 | **Average TTP (days)** | 60 |
| ***p*-value** | 0.0004 | ***p*-value** | 0.0669 |
| ***p*-value comparing lineages** | | 0.2275 | |
| **PDX PH039-A rucapraib 450 mg/kg response** | | **PDX PH039-B rucapraib 450 mg/kg response** | |
| **Response** | Responsive (SD) | **Response** | Responsive (CR) |
| **Median TTH (days)** | 60 | **Median TTH (days)** | 88 |
| **Average TTP (days)** | 50 | **Average TTP (days)** | 71 |
| ***p*-value** | 0.0003 | ***p*-value** | 0.0011 |
| ***p*-value comparing lineages** | | 0.0004 | |
| **PDX PH039-A niraparib 100 mg/kg response** | | **PDX PH039-B niraparib 100 mg/kg response** | |
| **Response** | Responsive (SD) | **Response** | Responsive (CR) |
| **Median TTH (days)** | 53 | **Median TTH (days)** | 71 |
| **Average TTP (days)** | 43 | **Average TTP (days)** | 64 |
| ***p*-value** | 0.0002 | ***p*-value** | 0.0508 |
| ***p*-value comparing lineages** | | 0.0766 | |

**Supplementary Table 4. PH039 Lineage B contains multiple tumors with "shifted heterogeneous" me*RAD51C* profile.** MS-HRM me*RAD51C* profiles for vehicle-treated PDX aliquots from PH039 lineages A and B, where "right-shifted heterogeneous" represents heterogeneous MS-HRM curves are shifted towards to the 100% methylated control curve (i.e. slightly more methylated).

| PH039 Lineage | PDX aliquot | Transplant | me*RAD51C* profile |
|---|---|---|---|
| **A** | 12970 | T4 | Heterogeneous |
| **A** | 12971 | T4 | Heterogeneous |
| **A** | 13039 | T4 | Heterogeneous |
| **A** | 13288 | T5 | Heterogeneous |
| **A** | 14002 | T7 | Heterogeneous |
| **A** | 14179 | T8 | Heterogeneous |
| **B** | 13070 | T4 | Heterogeneous |
| **B** | 13071 | T4 | Right-shifted heterogeneous |
| **B** | 13368 | T5 | Right-shifted heterogeneous |
| **B** | 13381 | T5 | Right-shifted heterogeneous |
| **B** | 13939 | T6 | Heterogeneous |
| **B** | 14224 | T7 | Heterogeneous |

**Supplementary Table 5. PDX #183 in vivo rucaparib re-treatment response summary.** TTH – time to harvest; TTP – time to progression.

| | |
|---|---|
| **PDX #183 vehicle values** | |
| **Median TTH (days)** | 50 |
| **Average TTP (days)** | 8 |
| **PDX #183 vehicle (following rucaparib cycle 2)** | |
| **Median TTH (days)** | 43 |
| **Average TTP (days)** | 8 |
| ***p*-value** | 0.475 |
| **PDX #183 rucaparib cycle 2** | |
| **Median TTH (days)** | 116 |
| **Average TTP (days)** | 47 |
| ***p*-value** | <0.0001 |
| **PDX #183 rucaparib cycle 4** | |
| **Median TTH (days)** | 99 |
| **Average TTP (days)** | 36 |
| ***p*-value** | <0.0001 |

**Supplementary Table 6. Summary of all results for me*RAD51C* loss screening by MS-HRM.** 164 tumor aliquots tested in total (across all PDX models).

| Treatment | Tumours with 0% me*RAD51C* | | Tumours with 1-50% me*RAD51C* | | Tumours with 51-99% me*RAD51C* | | Tumours with 100% me*RAD51C* | | Tumours with me*RAD51C* curve shift | |
|---|---|---|---|---|---|---|---|---|---|---|
| **PDX #183** | | | | | | | | | | |
| **Vehicle** | 0/21 | 0% | 0/21 | 0% | 0/21 | 0% | 0/21 | 0% | 1/21 | 4.8% |
| **Niraparib 100mg/kg** | 0/7 | 0% | 1/7 | 0% | 0/7 | 0% | 6/7 | 86% | 1/7 | 14.3% |
| **Rucaparib 300mg/kg** | 0/12 | 0% | 1/12 | 8% | 0/12 | 0% | 11/12 | 92% | 0/12 | 0.0% |
| **Rucaparib 450mg/kg** | 0/10 | 0% | 0/10 | 0% | 0/10 | 0% | 10/10 | 100% | 0/10 | 0.0% |
| **Rucaparib 300mg/kg cycle 2** | 0/7 | 0% | 0/7 | 0% | 0/7 | 0% | 7/7 | 100% | 0/7 | 0.0% |
| **Rucaparib 300mg/kg cycle 3** | 0/2 | 0% | 0/2 | 0% | 0/2 | 0% | 2/2 | 100% | 0/2 | 0.0% |
| **Rucaparib 300mg/kg cycle 4** | 0/11 | 0% | 0/11 | 0% | 3/11 | 27% | 8/11 | 73% | 0/11 | 0.0% |
| **Cisplatin 4mg/kg** | 0/7 | 0.0% | 0/7 | 0% | 1/7 | 14% | 6/7 | 86% | 0/7 | 0.0% |
| **PDX PH039-A** | | | | | | | | | | |
| **Vehicle** | 0/6 | 0% | 0/6 | 0% | 0/6 | 0% | 0/6 | 0% | 0/6 | 0.0% |
| **Niraparib 100mg/kg** | 0/4 | 0% | 1/4 | 25% | 0/4 | 0% | 0/4 | 0% | 0/4 | 0.0% |
| **Rucaparib 300mg/kg** | 0/7 | 0% | 2/7 | 29% | 2/7 | 29% | 3/7 | 43% | 1/7 | 14.3% |
| **Rucaparib 450mg/kg** | 1/6 | 17% | 2/6 | 33% | 0/6 | 0% | 3/6 | 50% | 0/6 | 0.0% |
| **Rucaparib 450mg/kg cycle 2** | 1/3 | 33% | 0/3 | 0% | 0/3 | 0% | 2/3 | 67% | 0/3 | 0.0% |
| **Rucaparib 450mg/kg cycle 3** | 3/3 | 100% | 0/3 | 0% | 0/3 | 0% | 0/3 | 0% | 0/3 | 0.0% |
| **PDX PH039-B** | | | | | | | | | | |
| **Vehicle** | 0/6 | 0% | 0/6 | 0% | 0/6 | 0% | 6/6 | 100% | 3/6 | 50.0% |
| **Niraparib 100mg/kg** | 0/4 | 0% | 0/4 | 0% | 0/4 | 0% | 4/4 | 100% | 1/4 | 25.0% |
| **Rucaparib 300mg/kg** | 0/4 | 0% | 0/4 | 0% | 0/4 | 0% | 4/4 | 100% | 0/4 | 0.0% |
| **Rucaparib 450mg/kg** | 1/5 | 20% | 2/5 | 40% | 0/5 | 0% | 2/5 | 40% | 0/5 | 0.0% |
| **Rucaparib 450mg/kg cycle 2** | 1/1 | 100% | 0/1 | 0% | 0/1 | 0% | 0/1 | 0% | 0/1 | 0.0% |
| **Rucaparib 450mg/kg cycle 3** | 5/5 | 100% | 0/5 | 0% | 0/5 | 0% | 0/5 | 0% | 0/5 | 0.0% |
| **Cisplatin 4mg/kg** | 0/3 | 0% | 0/3 | 0% | 0/3 | 0% | 0/3 | 100% | 3/3 | 0.0% |

**Supplementary Table 7.** Extended clinical information for the patient cohort. Samples grouped by me*RAD51C* pattern (homogeneous vs heterogeneous), and platinum TFI. Refractory defined as progressive disease on chemotherapy, resistant as PR or recurrence/progression within 6 months of end of chemotherapy and sensitive as recurrence/progression more than 6 months after end of chemotherapy. DOD – Died of disease; CR – complete response; PR – partial response; PD – progressive disease; SD – stable disease; N/A – Not available;  TFI – Treatment-Free Interval.

| **Patient ID** | **Surgical outcome** | **Lines of therapy received** | | | **First platinum TFI** | **Primary platinum status** | **Survival** | **Neoplastic cellularity (qpure %)** | ***RAD51C* copy number** | **Methylated alleles %** | **Adjusted me*RAD51C*** | **me*RAD51C* profile** | **Age at diagnosis** | **FIGO Staging** | **Histology** | **Neo-adjuvant chemo** | **Sample type** | **Disease course summary** |
|---|---|---|---|---|---|---|---|---|---|---|---|---|---|---|---|---|---|---|
| | | **Total** | **Platinum** | **Prior to sample collection** | | | | | | | | | | | | | | |
| **LS-158** | <5mm | 1 | 1 | 0 | 87 months | Sensitive | 87 months at last FU, alive | 90.26 | 2 | 93 | 100% | Homogeneous | 39 | IIIC | Serous Carcinoma | No | Primary tumor, primary surgery, snap-frozen | Sample taken at primary surgery. Adjuvant treatment with carboplatin / paclitaxel / topotecan (CR). Patient remains in first remission, with no evidence of disease at last follow-up. |
| **LS-215** | No residual disease | 3 | 3 | 0 | 22 months | Sensitive | 41 months, DOD | 56.43 | 2 | 72 | 100% | Homogeneous | 59 | IV | Poorly Differentiated | No | Primary tumor, primary surgery, snap-frozen | Sample taken at primary surgery. First line treatment with carboplatin / paclitaxel (CA125 normalization). Intra-abdominal disease and liver metastases after 18 months. Two additional lines of platinum chemotheray recieved, but patient chose to discontinue third line and died soon after. |
| **LS-267** | Suboptimal debulk | 3 | 1 | 0 | 18 months | Sensitive | 24 months, DOD | 58.93 | 2 | 75 | 100% | Homogeneous | 63 | IIIC | Serous Carcinoma | No | Primary tumor, primary surgery, snap-frozen | Sample taken at primary surgery. First line carboplatin / paclitaxel (CA125 normalization, negative CT), with monthly paclitaxel as maintenance therapy. Rising CA125 after 9 months, evidence of ascites and metastatic disease on CT. Monthly liposomal doxorubicin / bevacizumab 4 months (SD) until rising CA125. Gemcitabine / bevacizumab for 2 months (SD). Therapy discontinued (medical complications), patient died two months later. |
| **AOCS-106** | Macroscopic disease ≤1 cm | 2 | 1 | 1 | 4 months | Resistant (sample post-neoadjuvent chemotherpay) | 12 months, DOD | 92.14 | 3 | 55 | 58% | Homogeneous and heterozygous | 64 | IIIC | Serous Carcinoma | Yes | Primary tumor, interval debulk, snap frozen | Neoadjuvant carboplatin / paclitaxel, sample taken at primary surgery nine months later. Second line peglyated liposomal doxorubicin started 7 months after surgery. Patient died due to disease progression. |
| **LS-376** | No residual disease | 5 | 3 | 0 | 2 months | Resistant | 25 months, DOD | 95.56 | 3 | 89 | 92% | Homogeneous | 61 | IIIC | Serous Carcinoma | No | Primary tumor, primary surgery, snap-frozen | Sample taken at primary surgery. First line carboplatin / paclitaxel. Second line carboplatin/ gemcitabine / PARPi (veliparib). Ascites and elevated CA125 five and a half months post-chemotherapy. Third line topotecan/ bevacizumab (progressive ascites). Switched to nab-paclitaxel for two months. One cycle of carboplatin before dying two months later. |
| **LS-28** | Suboptimal debulk, >2cm | ≥1 | ≥1 | 0 | N/A | Refractory | 25 months, DOD | N/A | N/A | 72 | N/A | Homogeneous | 54 | IIIC | Serous Carcinoma | No | Primary tumor, primary surgery, snap-frozen | Sample taken at primary surgery. First line carboplatin/ paclitaxel (CA125 normalization) . Six months post-surgery small positive foci detected in periaortic lymph nodes. Second line topotecan (PD). No further treatment records available until her death 19 months after second surgery. |
| **AOCS-120** | Macroscopic disease ≤1 cm | 6 | 4 | 6 | 17 months | Sensitive (sample from resistant recurrence) | 76 months, DOD | 92.87 | 3 | 96 | 100% | Heterogeneous | 46 | IIIC | Serous Carcinoma | No | Resistant recurrence, ascites, snap frozen | First line carboplatin / paclitaxel / gemcitabine. 17 months later second line carboplatin / paclitaxel. 20 months later third line carboplatin / gemcitabine. Six months later fourth line carboplatin. Nine months later fifth line pegylated liposomal doxorubicin. Three months later, another line (6th) of pegylated liposomal doxorubicin. Sample taken one month after end of final therapy. Death due to disease progression. |
| **AOCS-143** | Macroscopic disease ≤1 cm | 9 | 6 | 0 | 7 months | Sensitive | 69 months, DOD | 78.30 | 2 | 81 | 100% | Heterogeneous | 49 | IIIC | Serous Carcinoma | No | Primary tumour, primary surgery, snap-frozen | Sample taken at primary surgery. First line carboplatin / paclitaxel, second line pegylated liposomal doxorubicin, third line carboplatin / gemcitabine. Tamoxifen. Fourth line docetaxel. Fifth line carboplatin / docetaxel / etoposide. One month later sixth line cisplatin / etoposide. Four months later seventh line cisplatin. Five months later eight line paclitaxel. One month later ninth line cisplatin. Death due to acute leukaemia related to chemotherapy. |
| **LS-473** | Suboptimal debulk, 3-4cm | 6 | 3 | 1 | 6 months | Resistant (sample post-neoadjuvent chemotherpay) | 45 months, DOD | 79.43 | 4 | 86 | 97% | Heterogeneous | 51 | IIIC | Serous Carcinoma | Yes | Primary tumor, primary surgery, snap-frozen | Neoadjuvant carboplatin / paclitaxel (PR). Sample obtained at interval debulking surgery. Carboplatin / paclitaxel / bevacizumab post-surgery. Complete metabolic response (PET scan). Rising CA125 and PD just prior to six months post chemotherapy, thus re-commenced carboplatin / paclitaxel / bevacizumab. Rising CA125 and PD after seven months. Third line gemcitabine (rising CA125 and abdominal symptoms after two months). Gemcitabine / bevacizumab for three months (rising CA125 and PD). Liposomal doxorubicin started but switched to cisplatin / pemetrexed after three months (rising CA125 and small bowel obstruction). Lost to follow up until death nine months later. |
| **LS-472** | Suboptimal debulk, 1.5cm | 4 | 1 | 0 | N/A | Resistant | 26 months, DOD | 97.36 | 1 | 94 | 99% | Heterogeneous | 54 | IIIC | Serous Carcinoma | No | Primary tumor, primary surgery, snap-frozen | Sample taken at primary surgery. First line dose-dense paclitaxel/ carboplatin (normalization of CA125), but persistent disease. Additional three lines of chemotherapy (bevacizumab, topotecan, Doxil) with PD on all. |
| **LS-467** | No residual disease | N/A | N/A | 0 | N/A | N/A | N/A | 75.91 | 3 | 88 | 100% | Heterogeneous | 67 | IIB | Serous & Clear Cell | No | Primary tumor, primary surgery, snap-frozen | Sample taken at primary surgery. Patient lost to follow-up from commencement of first-line chemotherapy (carboplatin/ paclitaxel, one month after surgery). |
| **SFRC-1307** | Suboptimal debulk | 0 | 0 | 0 | N/A | N/A | 3 months at last FU, alive | N/A | N/A | N/A | N/A | Heterogeneous* | 45 | IIIC | Serous Carcinoma | No | Primary tumor, primary surgery, snap-frozen | Sample taken at primary surgery. Patient yet to begin treatment plan of adjuvant chemotherapy / bevacizumab. |

**Supplementary Table 8. Adjusted me*RAD51C* for two pieces of tumor LS-473**

| Patient LS-473 | Neoplastic cellularity (%) | *RAD51C* copy number | Methylated alleles % | Adjusted me*RAD51C* | me*RAD51C* profile |
|---|---|---|---|---|---|
| **Piece 1** | 79 | 4 | 86 | 97% | Heterogeneous |
| **Piece 2** | 89 | 4 | 80 | 85% | Heterogeneous |

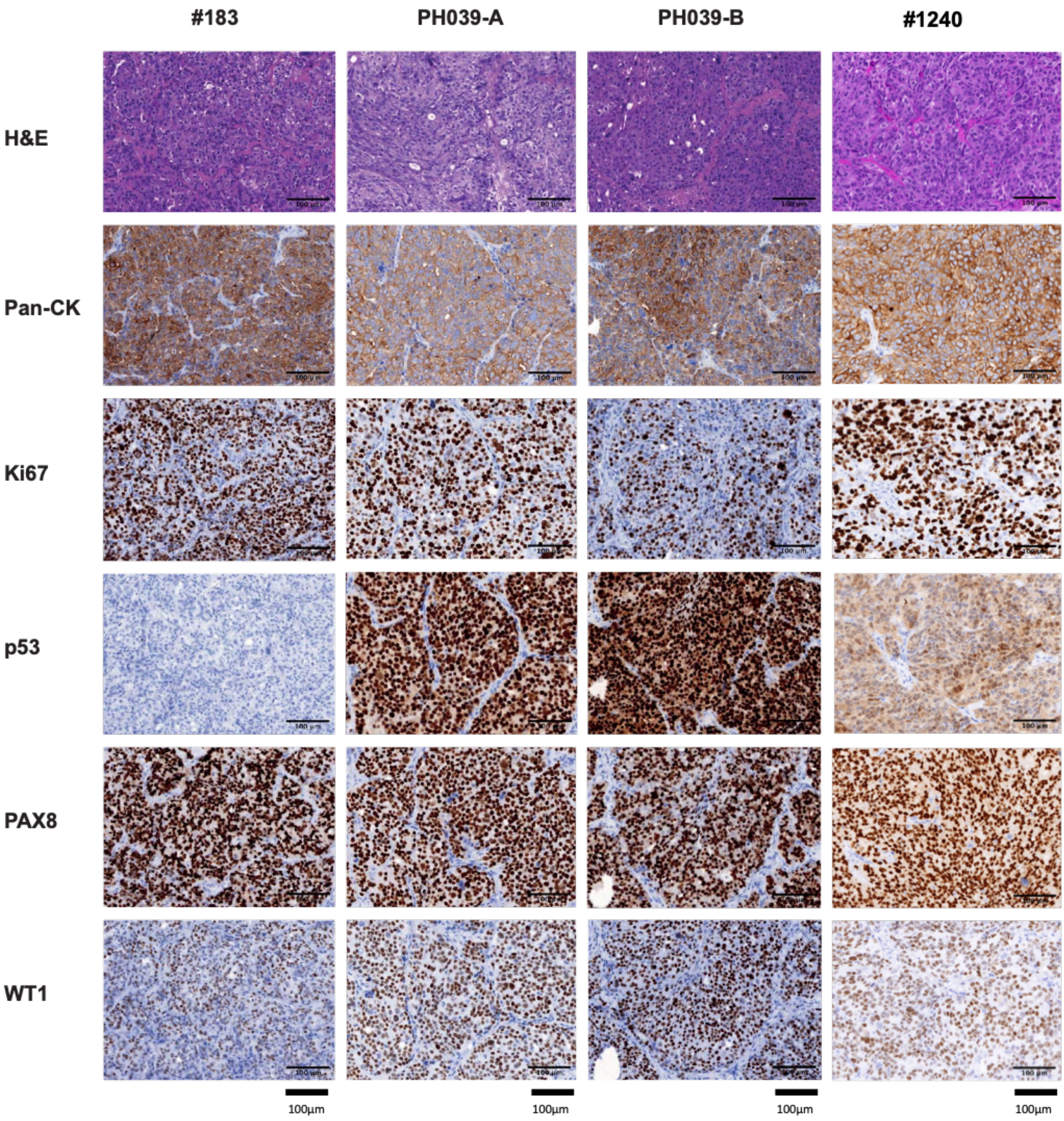


**Supplementary Figure 1. Histology of each PDX model.** Immunohistochemistry (IHC) staining is consistent with HGSC histology for both models (high PAX8). TP53 staining is absent in #183 as expected (nonsense *TP53* mutation NM_000546.6 :c.661G>T (p.E221*)). The two lineages of PH039 (PH039-A and PH039-B) have consistent staining and share morphological appearance. Scale bar is 100um.

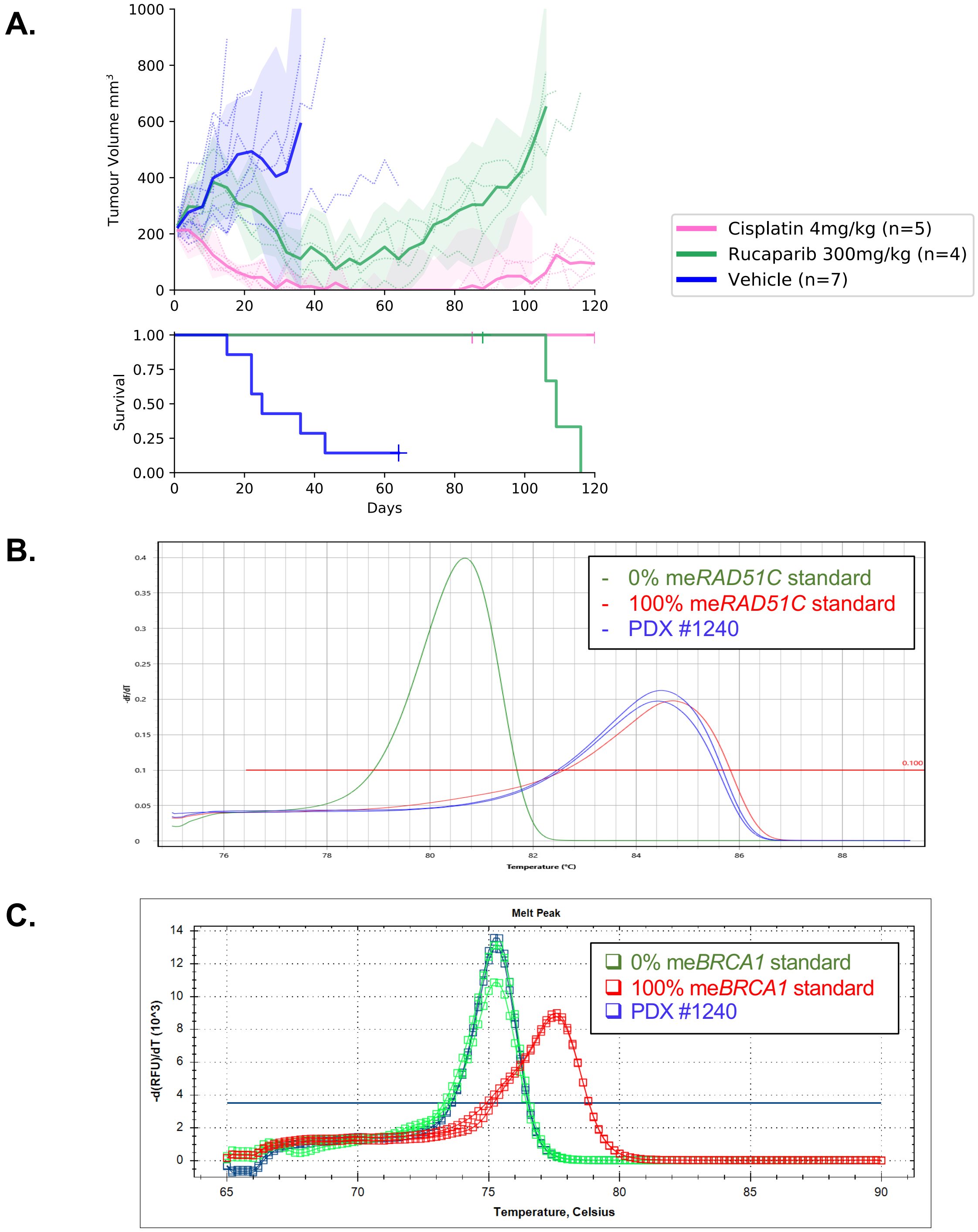


**Supplementary Figure 2. Characterization of PDX #1240. (A)** *In vivo* responses of PDX #1240 to cisplatin (4mg/kg) and rucaparib (300mg/kg). Mean tumor volume ($mm^3$) ± 95% CI (hashed lines are representing individual mice) and corresponding Kaplan–Meier survival analysis. Censored events are represented by crosses on Kaplan–Meier plot; n = individual mice. MS-HRM results for **(B)** me*RAD51C* (MIC platform) and **(C)** me*BRCA1* (CFX platform) in an untreated T1 PDX #1240 aliquot.

A.

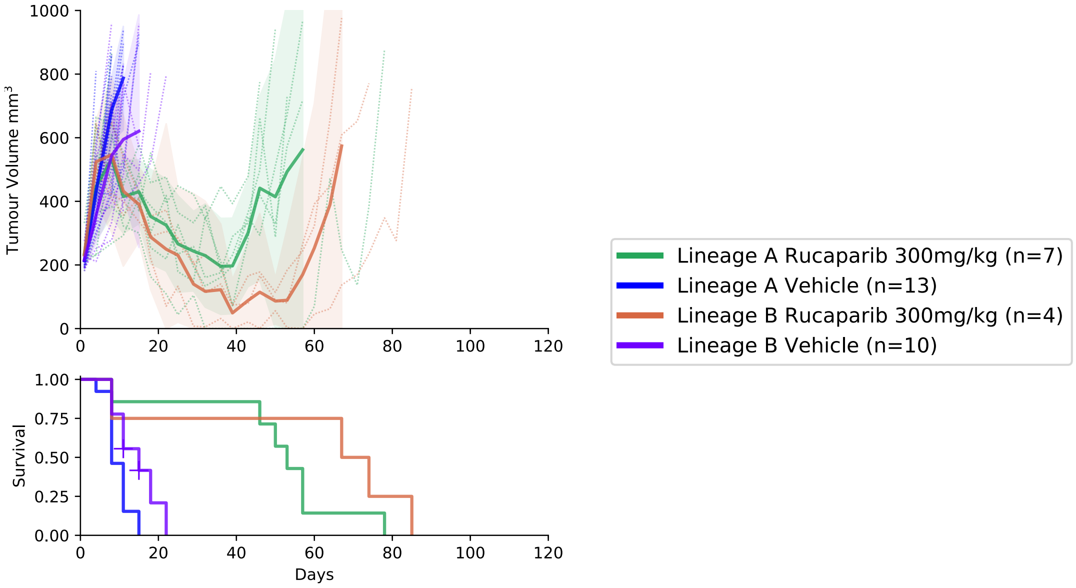


B.

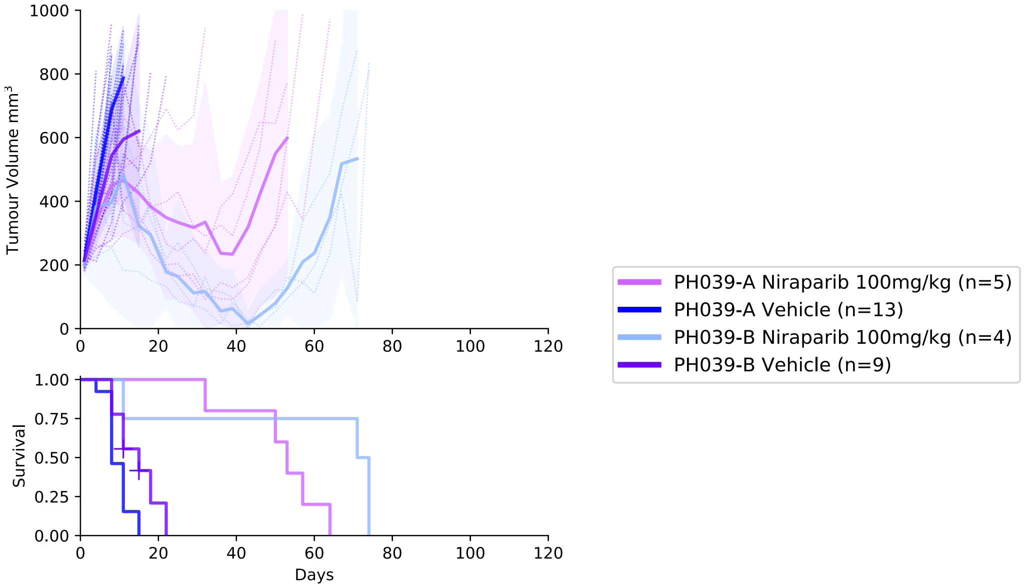


**Supplementary Figure 3. WEHI-PH039 Lineage A vs B *in vivo* PARPi responses.** PDX WEHI-PH039 Lineages A and B are both show some response to **(A)** rucaparib (300 mg/kg) and **(B)** niraparib (100 mg/kg), though lineage B was consistently more responsive than lineage A. Summary statistics presented in Supplementary Table 3. Mean tumor volume ($mm^3$) ± 95% CI (hashed lines are representing individual mice) and corresponding Kaplan–Meier survival analysis. Censored events are represented by crosses on Kaplan–Meier plot; n = individual mice.

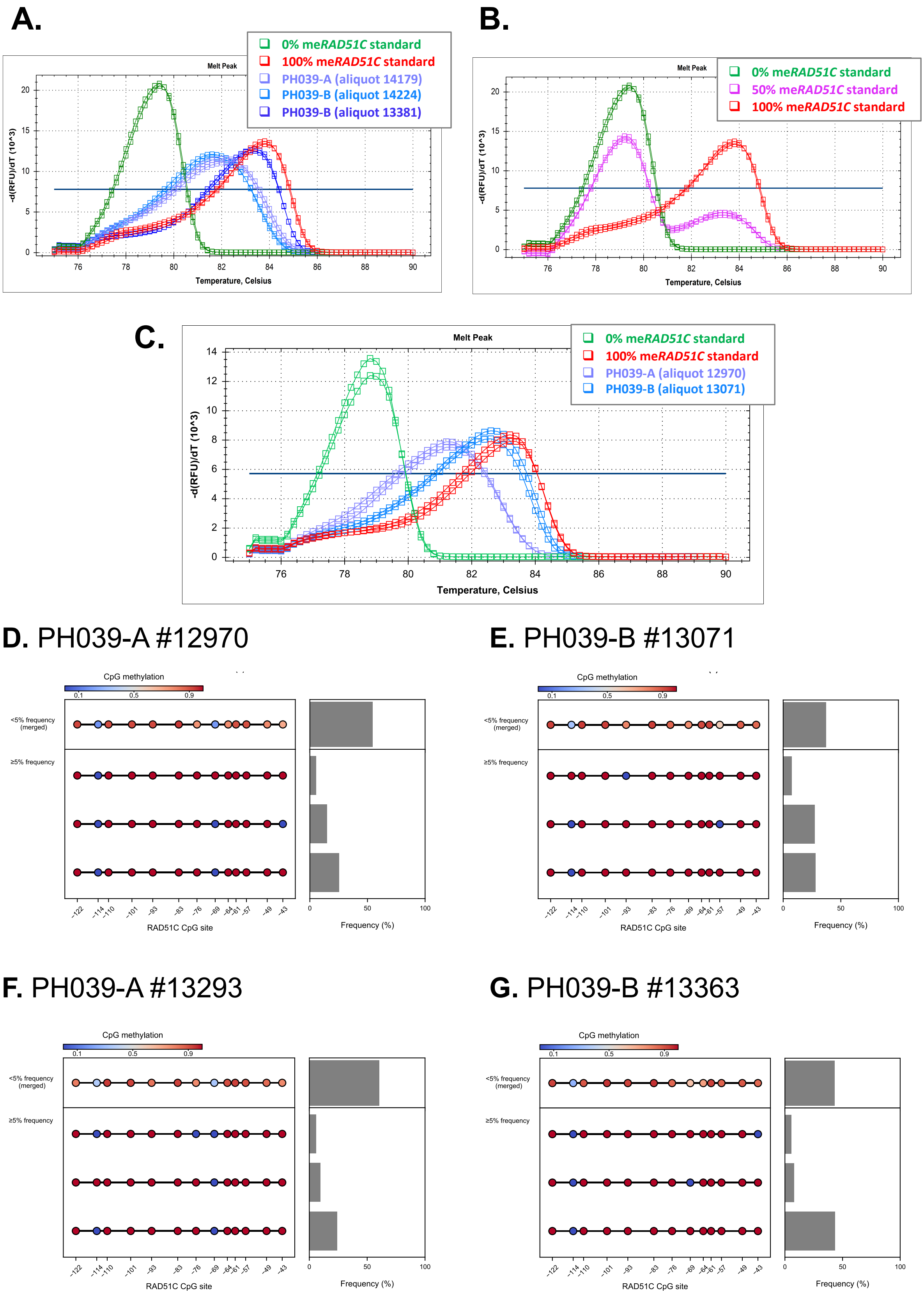

A.
Melt Peak
0% meRAD51C standard
100% meRAD51C standard
PH039-A (aliquot 14179)
PH039-B (aliquot 14224)
PH039-B (aliquot 13381)
-d(RFU)/dT (10^3)
Temperature, Celsius
B.
Melt Peak
0% meRAD51C standard
50% meRAD51C standard
100% meRAD51C standard
-d(RFU)/dT (10^3)
Temperature, Celsius
C.
Melt Peak
0% meRAD51C standard
100% meRAD51C standard
PH039-A (aliquot 12970)
PH039-B (aliquot 13071)
-d(RFU)/dT (10^3)
Temperature, Celsius
D. PH039-A #12970
E. PH039-B #13071
F. PH039-A #13293
G. PH039-B #13363
CpG methylation
0.1
0.5
0.9
<5% frequency (merged)
≥5% frequency
−122
−114
−110
−101
−93
−83
−76
−69
−64
−61
−57
−49
−43
RAD51C CpG site
Frequency (%)

**Supplementary Figure 4. PDX PH039-B contains aliquots with either heterogeneous and "right-shifted heterogeneous" me*RAD51C* profiles. (A)** Example MS-HRM profiles of PH039-A and PH039B aliquots (14179 and 14224) with heterogeneous me*RAD51C*, and a PH039-B aliquot (13381) with "right-shifted heterogeneous" me*RAD51C*. **(B)** Example of heterozygous me*RAD51C* profile from 50% me*RAD51C* standard. **(C)** MS-HRM curves of the PH039-A and PH039-B tumor aliquots chosen for sequencing (same bisulfite-converted DNA samples used for MS-HRM and NGS to allow direct comparison between the two techniques). **(D)** Targeted sequencing results for PH039-A aliquot 12970 ("heterogeneous" MS-HRM profile) show two CpG sites (-114 and -69) are unmethylated in most epialleles. **(E)** Targeted sequencing results for PH039-B aliquot 13071 ("right-shifted heterogeneous" MS-HRM profile) shown only CpG site -114 is unmethylated in most epialleles. **(F)** Targeted sequencing results for rucaparib 300 mg/kg-treated PH039-A aliquot 13293 ("heterogeneous" MS-HRM profile) show two CpG sites (-114 and -69) are unmethylated in most epialleles. **(G)** Targeted sequencing results for rucaparib 300 mg/kg-treated PH039-B aliquot 13363 ("right-shifted heterogeneous" MS-HRM profile) shown only CpG site -114 is unmethylated in most epialleles. This is consistent with the profiles found in vehicle-treated aliquots.

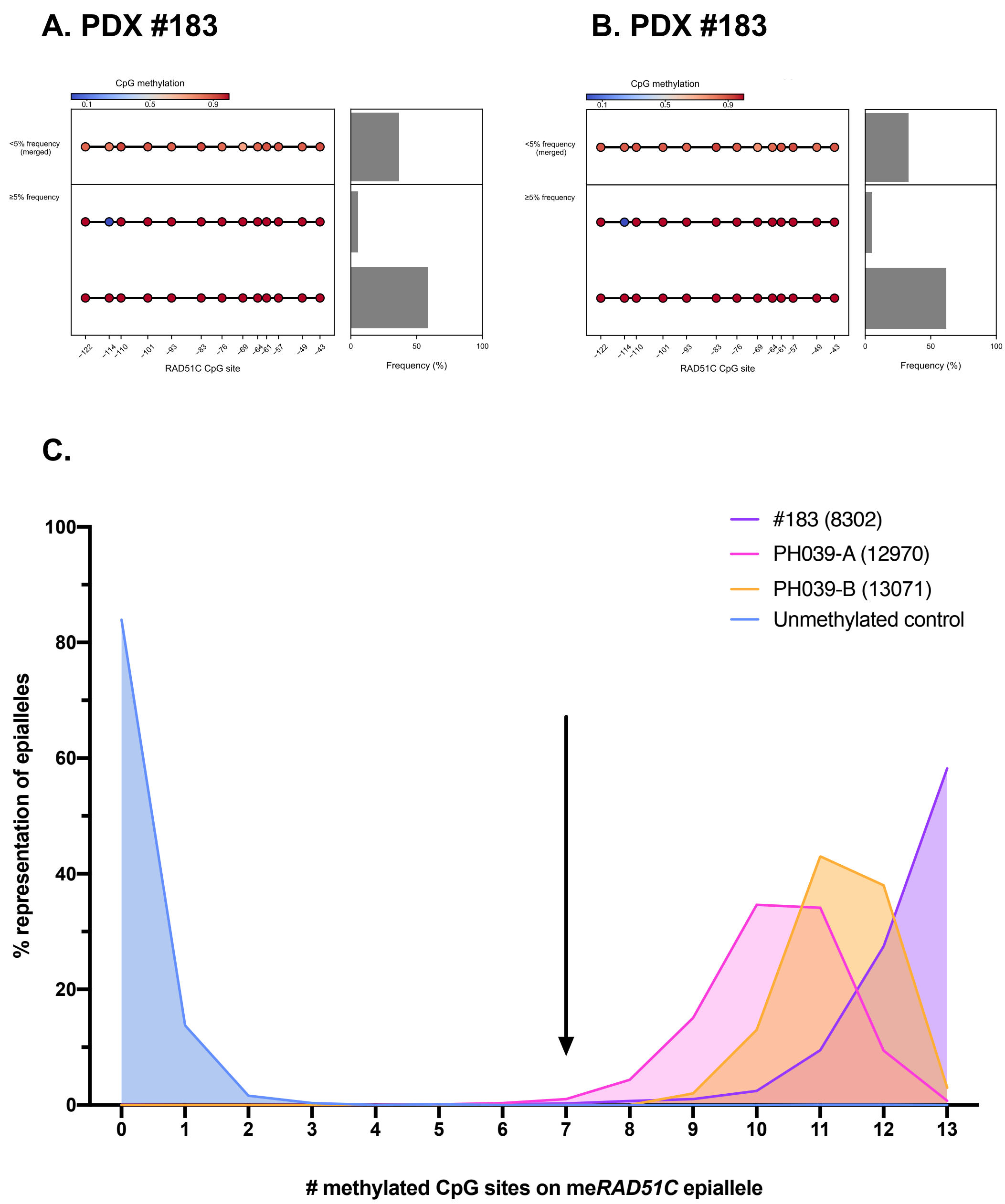


**Supplementary Figure 5. Epiallele distributions of PDX PH039 and #183 from targeted sequencing.** Sample me*RAD51C* sequencing profiles of vehicle-treated PDX #183 aliquots **(A)** 8302 and **(B)** 9423 reveal a high degree of epiallelic homogeneity compared to the PDX PH039-A and PH039-B tumor aliquots presented in Supplementary Fig. 5. **(C)** PH039-A tumor aliquots contain a minimum of 8 methylated CpG sites per epiallele (~60% of region methylated). PH039-B samples generally contain epialleles with the same number of methylated CpG sites, or more. Given tumors with these me*RAD51C* profiles are associated with *RAD51C* gene silencing, it appears that >60% methylation across this region is sufficient for gene silencing.

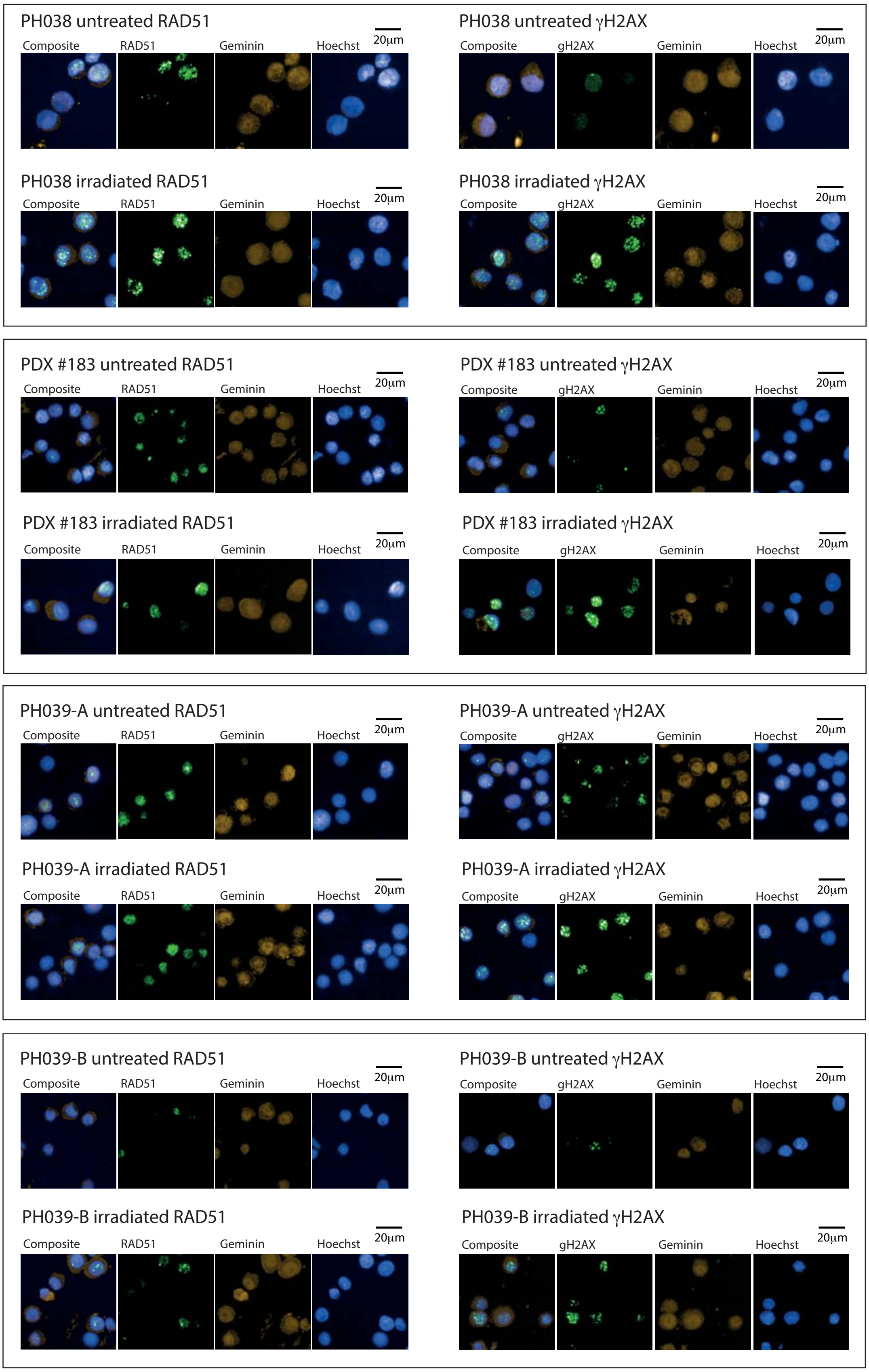


**Supplementary Figure 6. Representative RAD51 and γH2AX foci images from HRC control PDX PH038, and me*RAD51C* PDX models #183, PH039-A and PH039-B.**

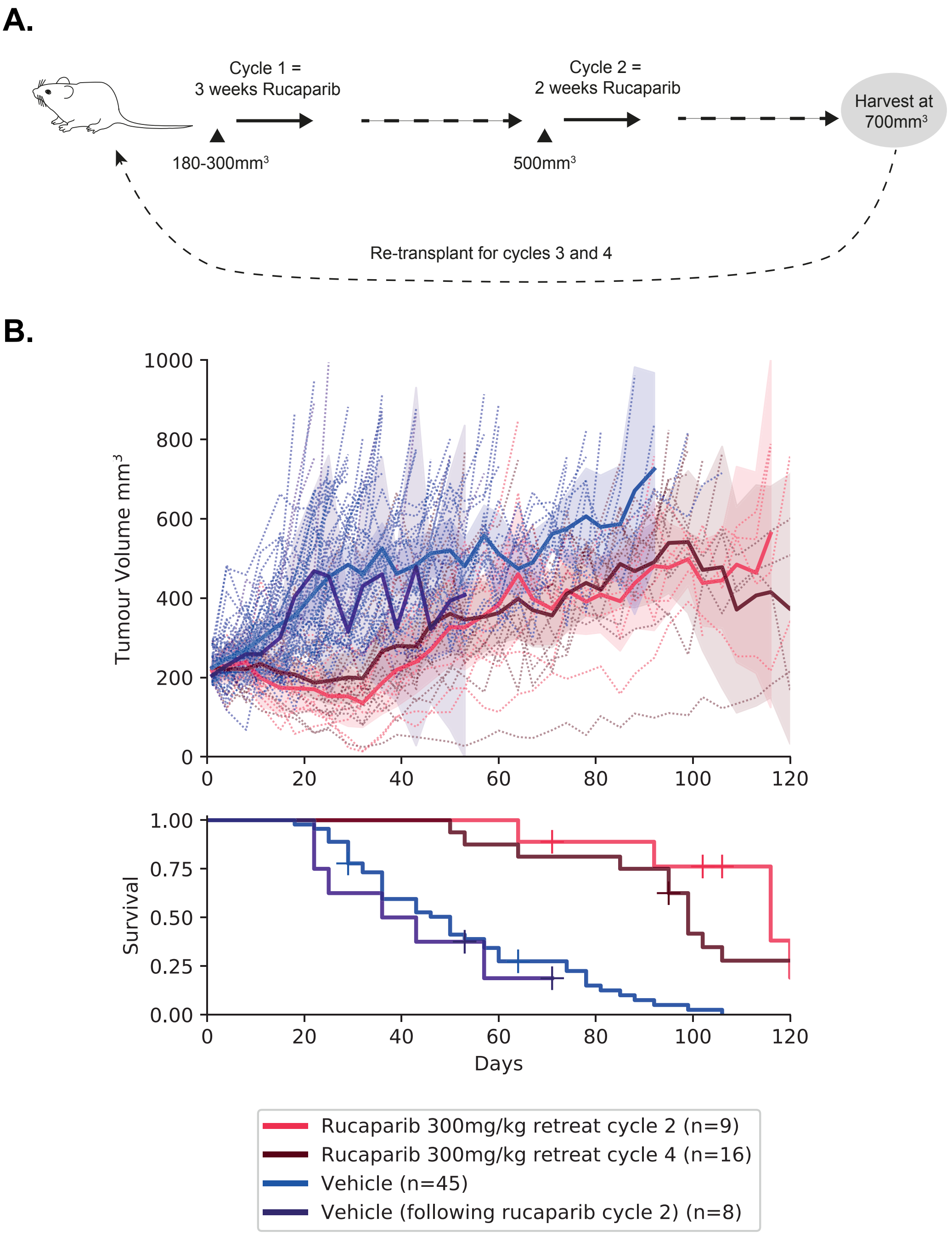


**Supplementary Figure 7. PDX #183 *in vivo* rucaparib re-treatment data. (A)** Rucaparib 300 mg/kg retreatment strategy used for PDX #183. **(B)** No difference in rucaparib response was observed for the cycle 2 vs cycle 4 tumor aliquots (p=0.2751). Mean tumor volume (mm3) ± 95% CI (hashed lines are representing individual mice) and corresponding Kaplan–Meier survival analysis. Censored events are represented by crosses on Kaplan–Meier plot; n = individual mice

**A.** #183 aliquot #13099 MS-HRM

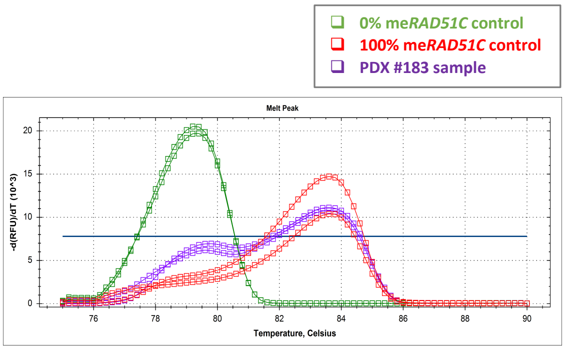


**B.** #183 aliquot #13099 NGS

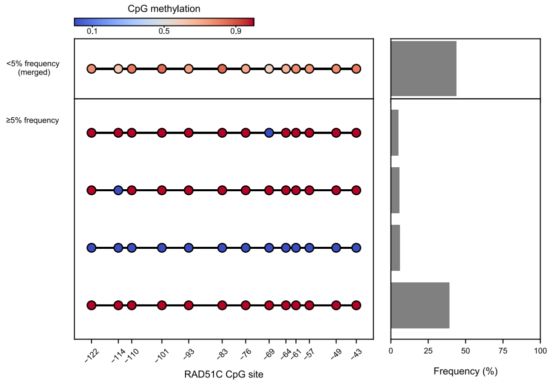


**C.** #183 aliquot 13393 MS-HRM

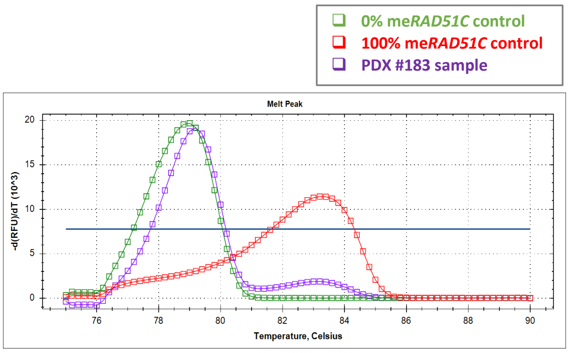


**D.** #183 aliquot 13393 NGS

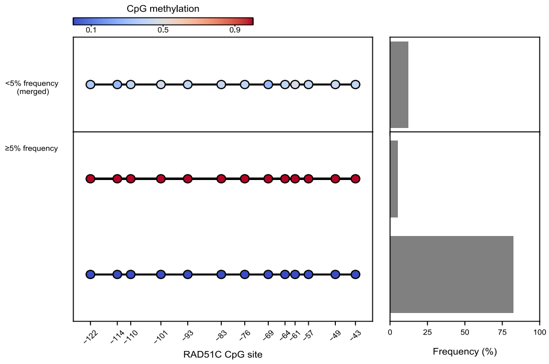


**E.**

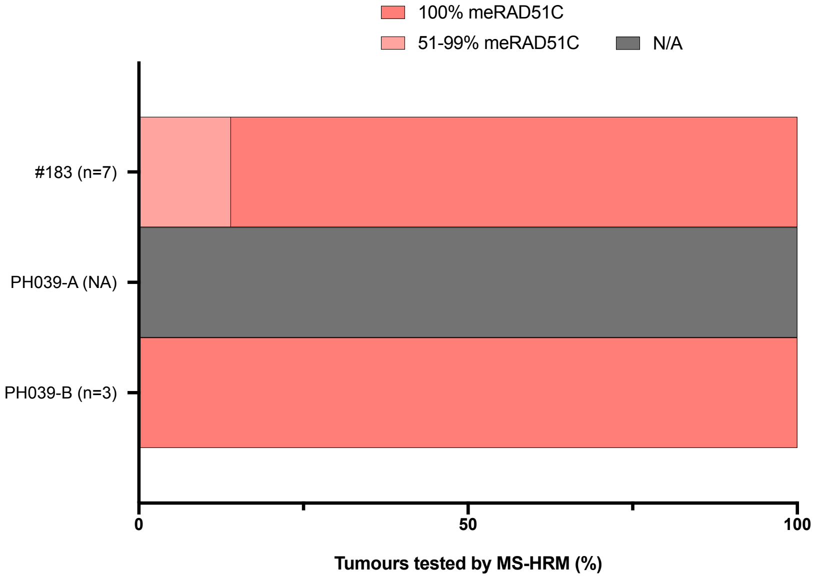

**Supplementary Figure 8. Loss of me*RAD51C* in PDX #183 samples. (A)** PDX #183 aliquot 13099 had estimated ~10-20% me*RAD51C* loss by MS-HRM following four cycles of rucaparib 300 mg/kg, and **(B)** this was found to be exactly 6% by targeted me*RAD51C* bisulfited sequencing (NGS). **(C)** PDX #183 aliquot 13393 had estimated ~90% me*RAD51C* loss by MS-HRM, **(D)** and was found to be exactly 83% by targeted me*RAD51C* bisulfited sequencing (NGS), with 5% only methylated alleles remaining. Each MS-HRM line represents a measurements per PDX aliquot/sample. RFU – Relative fluorescence units, Y axis is the derivative of fluorescence over temperature (-d(RFU)/dT) versus temperature (T). **(E)** PDX #183 (n=7) had some me*RAD51C* loss in one PDX aliquot following cisplatin, while PH039-B (n=3) did not. PH039-A was too sensitive to cisplatin and no PDX material was available for this lineage (N/A – not available).

**A.** PH039-A

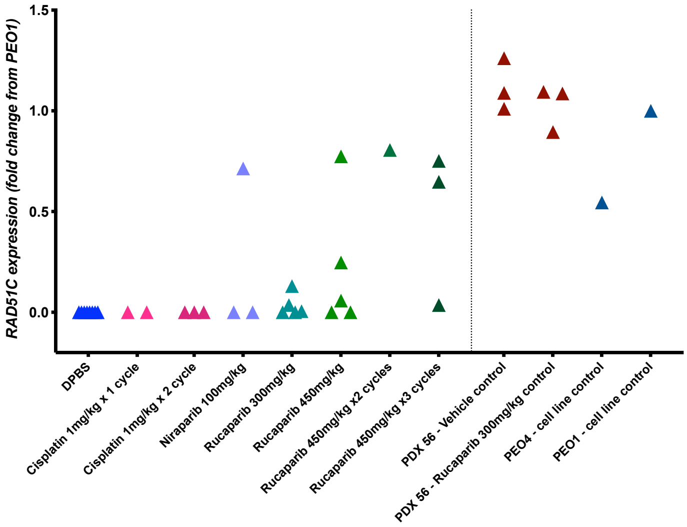


**B.** PH039-B

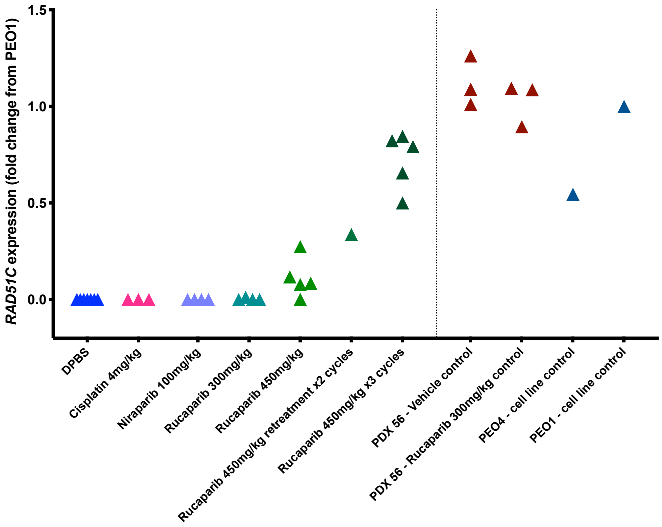


**C.** #183

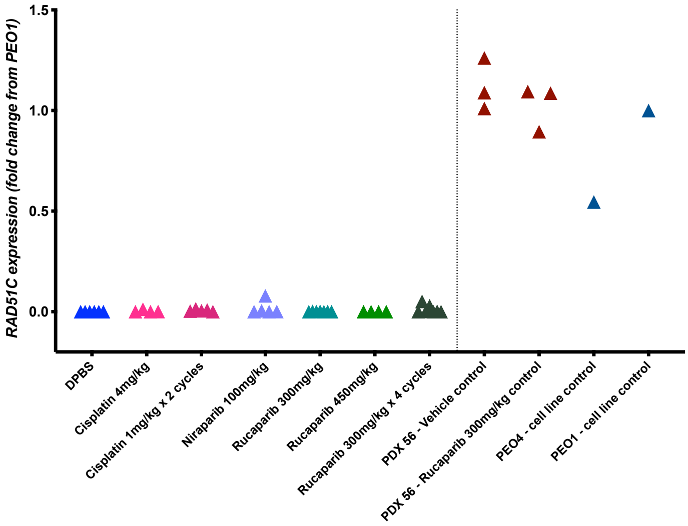


**D.**

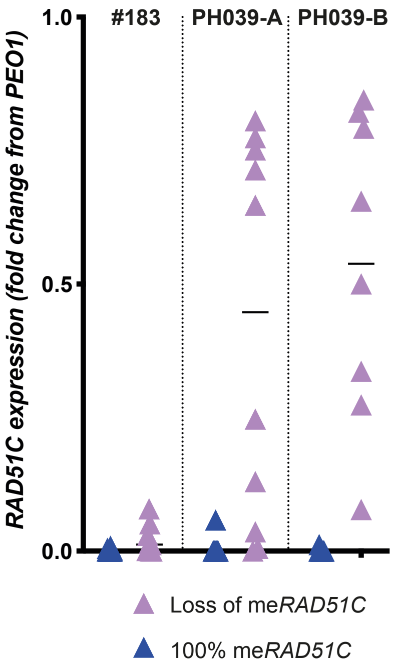

**Supplementary Figure 9. PDX *RAD51C* gene expression by qRT-PCR.** Each point represents an individual PDX tumor aliquot, presented as fold-change *RAD51C* expression from *BRCA2*-mutant cell line PEO1. **(A)** Results for all PDX PH039-A tumor aliquots, **(B)** all PDX PH039-B tumor aliquots and **(C)** all PDX #183 tumor aliquots tested for *RAD51C* gene expression by qRT-PCR. (D) Presented is the same data in A-C grouped by me*RAD51C* status (for samples where this was tested). Samples with any degree of me*RAD51C* loss (on MS-HRM) had significantly elevated *RAD51C* gene expression compared to samples with 100% me*RAD51C* (i.e. no me*RAD51C* loss detected) across all PDX models (data presented as fold-change from PEO1; $p<0.0001$ for all models using Mann-Whitney two-tailed test).

**A.**

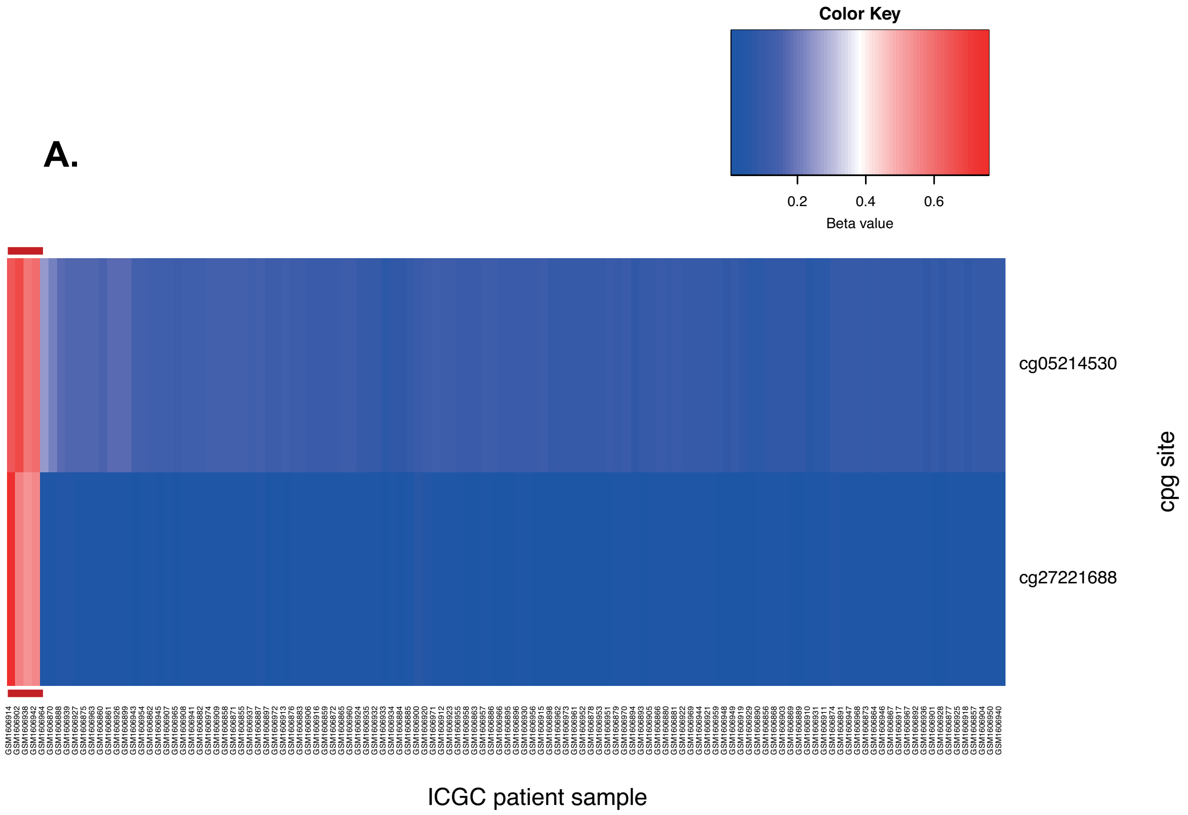


**B.**

| CpG | AOCS-106 | AOCS-120 | AOCS-143 | AOCS-147 |
|---|---|---|---|---|
| cg05214530 | 0.67 | 0.65 | 0.59 | 0.61 |
| cg27221688 | 0.57 | 0.76 | 0.54 | 0.56 |

**C.**

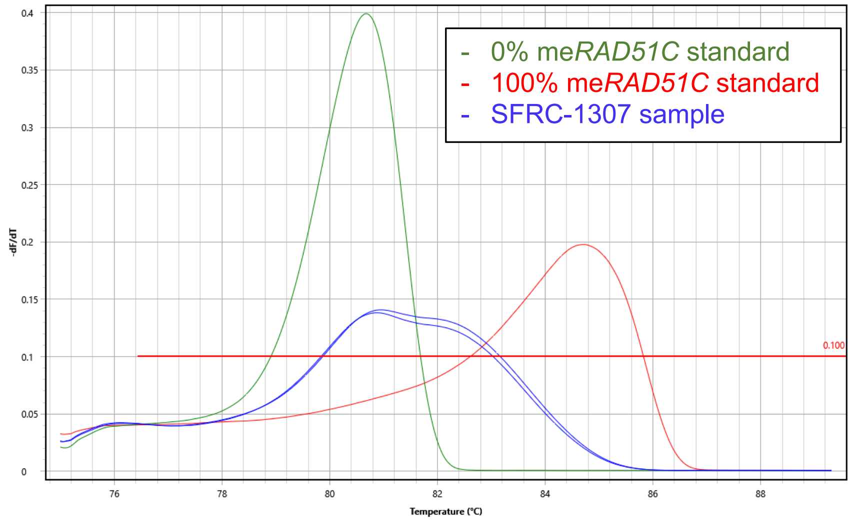

**Supplementary Figure 10. Identification of HGSC samples with *meRAD51C* in ICGC and SFRC cohorts. (A)** Four AOCS/Westmead GynBiobank patient samples from the ICGC-Ovarian Cancer Project (of 121 total samples) were found to have me*RAD51C* using publicly available Illumina 450K methylation array data from ICGC. The two CpG sites interrogated were covered by MS-HRM and targeted me*RAD51C* sequencing assays (cg05214530 is NM_058216.2:c.-114/ chr17:g.56769891 and cg27221688 is NM_058216.2:c.-101/ chr17:g.56769904). Samples were considered to have me*RAD51C* if CpG sites had a beta value of >0.5. Red bar indicates the four me*RAD51C* cases that were identified in this analysis. **(B)** Table of beta values for each CpG site within each AOCS sample with me*RAD51C*. **(C)** Recently diagnosed HGSC case SFRC-1307 was found to have me*RAD51C* in omental tumor sample by MS-HRM (MIC platform).

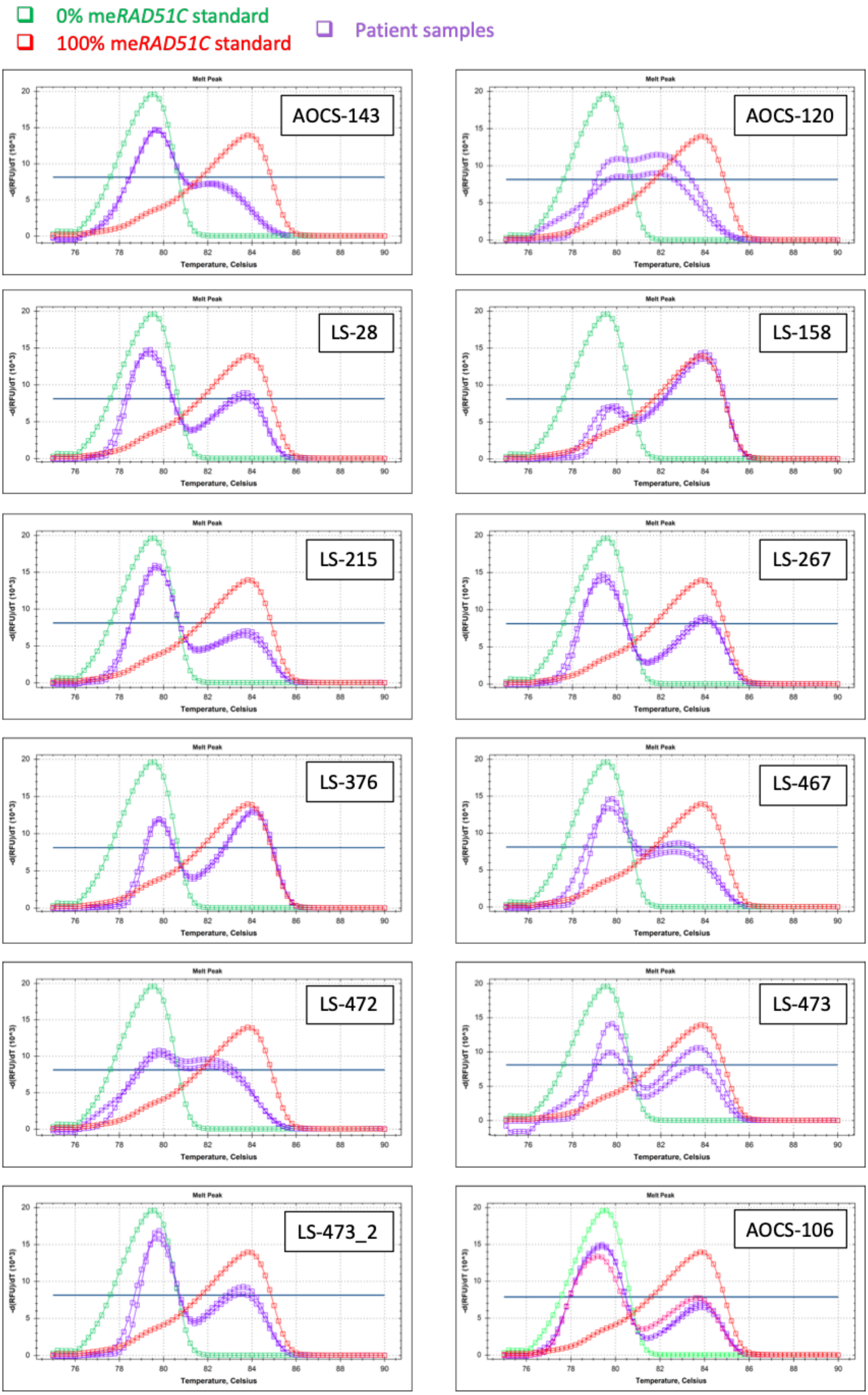


**Supplementary Figure 11. MS-HRM me*RAD51C* results for AOCS and LS patient samples.**

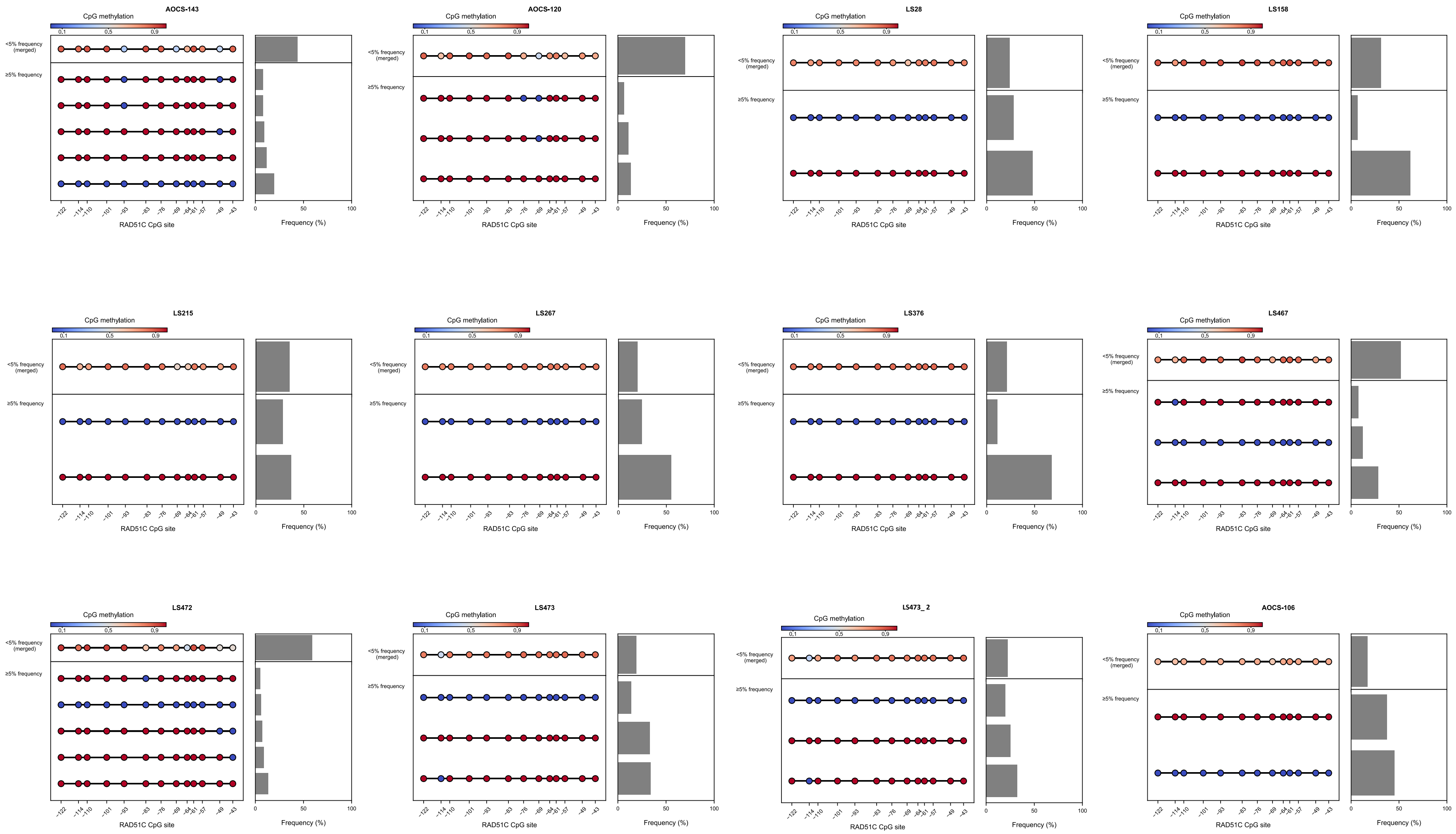


**Supplementary Figure 12. Targeted me*RAD51C* sequencing results for AOCS and LS patient samples.**

### Supp. Figure 13

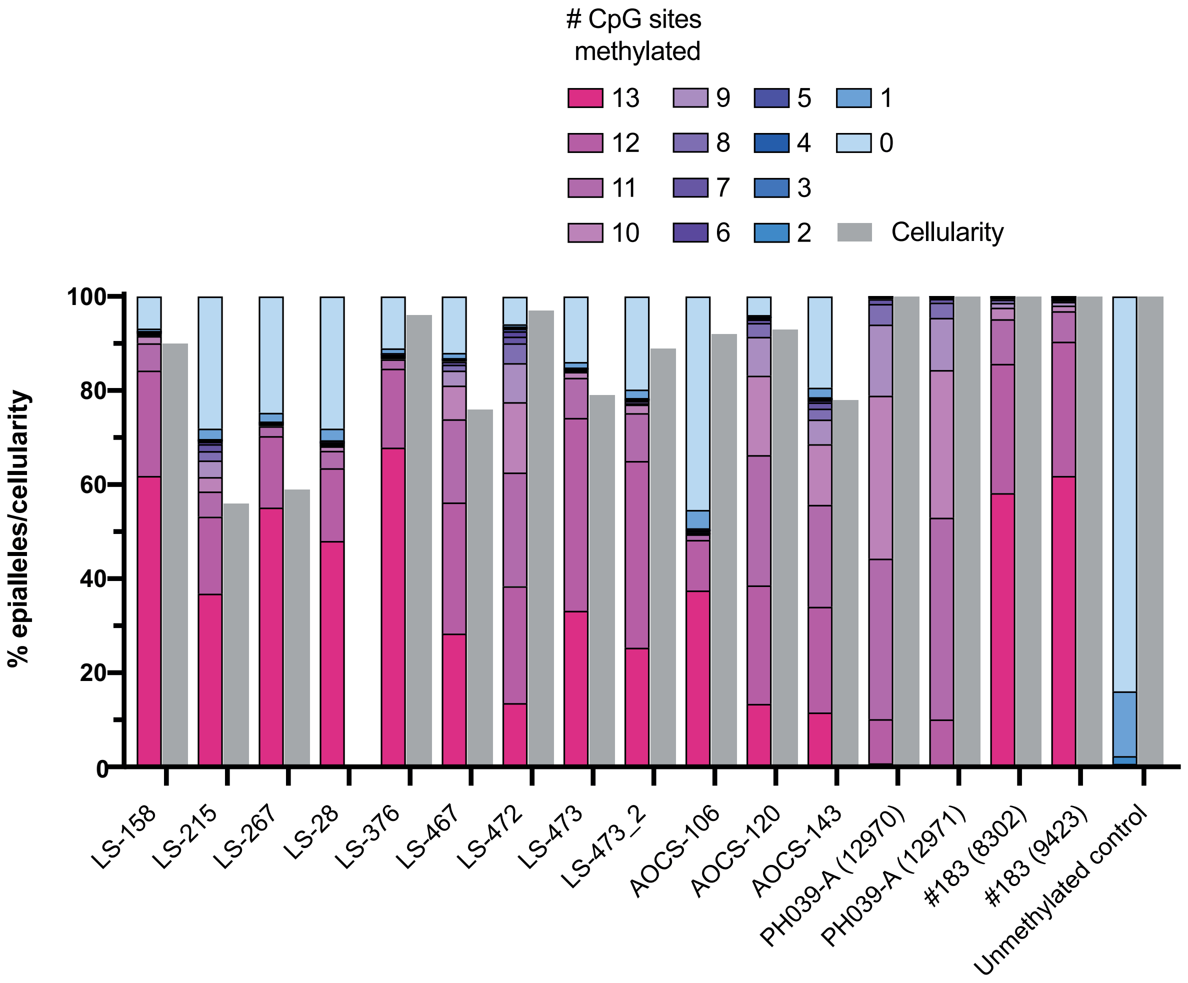


**Supplementary Figure 13. Grouping of epialleles by degree of CpG methylation in HGSC samples.** Presented are the targeted me*RAD51C* sequencing profiles of patient and PDX samples based on the representation of epialleles with varying degrees of CpG methylation (number of CpG sites methylated). Estimated cellularity based on SNP arrays is also included for reference. Unmethylated control is aliquot of PDX #62 (*BRCA1*-methylated, *RAD51C*-unmethylated).

### Supp. Figure 14

**A.**

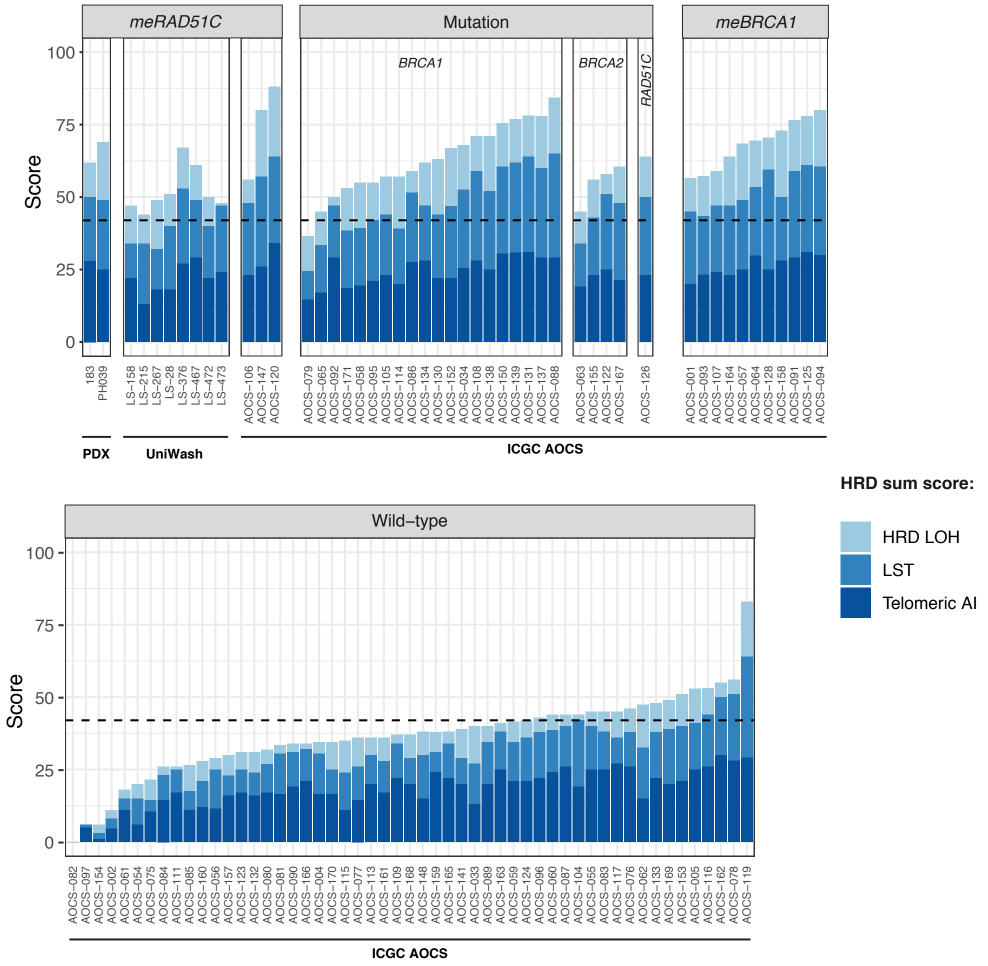


**B.**

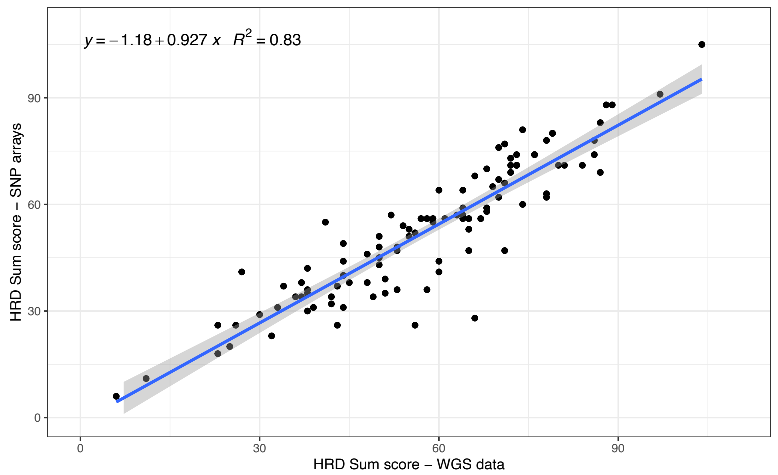

**Supplementary Figure 14. SNP array HRD scores for PDX and patient samples. (A)** All me*RAD51C* HGSC tested on SNP arrays in this study, including patient and PDX samples, were found to have high HRD scores, indicating either current or historical HRD in the tumor. SNP arrays of AOCS/ GynBiobank samples were performed previously in the ICGC-Ovarian Cancer Project[30] and were re-analyzed for this study. **(B)** There is a high correlation for HRD scores when comparing results from SNP arrays vs WGS ($R^2 = 0.83$), indicating that the SNP array data is reliable for estimating HRD score.
